# IMMF: An Interpretable Multi-Modal Framework for Hypothesis-Driven Biomarker Discovery in Triple-Negative Breast Cancer Using Public Data

**DOI:** 10.64898/2026.08.19.745809

**Authors:** AL Imran, Khandokar Md. Rahat Hossain, SM Rafiqul Islam, M. Sohel Rahman

## Abstract

Triple-Negative Breast Cancer (TNBC) is characterized by high heterogeneity, poor prognosis, and limited targeted treatment options. Bridging the gap between molecular alterations and histopathological morphology remains a major challenge in precision oncology. We propose an interpretable, multi-modal framework that integrates histopathological image analysis with multi-omics profiling (somatic mutations, DNA methylation, copy number alterations), leveraging U-Net-based nuclei segmentation, vision-language models (BLIP), biomedical language models (BioGPT), and explainable AI (SHAP, LIME). Our framework achieves strong predictive performance (AUC = 0.989) and provides transparent, biologically grounded interpretations by integrating morphological features with genomically prioritized biomarkers. Cross-modal analysis confirms established TNBC drivers and generates novel, testable hypotheses associating specific epigenetic alterations with distinct morphological phenotypes. While causal validation requires future wet-lab experiments, our framework accelerates hypothesis-driven biomarker discovery by integrating complementary data modalities with language-based reasoning, providing a transparent foundation for hypothesis generation and clinical translation.

## I. Introduction

Triple-Negative Breast Cancer (TNBC) is one of the most aggressive and clinically complex forms of breast cancer, accounting for roughly 15-20% of all diagnosed cases world-wide [1]. Defined by the absence of estrogen receptor (ER), progesterone receptor (PR), and human epidermal growth factor receptor 2 (HER2), TNBC lacks established molecular targets and remains resistant to endocrine or HER2-directed therapies [2]. Consequently, patients with TNBC experience early recurrence, rapid metastatic progression, and poorer overall survival than patients with hormone receptor-positive or HER2-enriched subtypes [3]. This absence of ER, PR, and HER2 expression is precisely what leaves TNBC resistant to targeted hormonal or HER2-directed therapies and underlies its comparatively poor prognosis relative to other breast cancer subtypes [1], [3].

Despite major advances in genomic profiling, TNBC remains highly heterogeneous at both the molecular and morphological levels. Large-scale genomic studies by Lehmann *et al*. [2] and Curtis *et al*. [4] established that TNBC is not a single disease but a spectrum of molecular subtypes including basal-like, mesenchymal, and immunomodulatory forms while Pereira *et al*. [5] further linked TNBC subtypes to specific somatic mutation and copy-number alteration patterns. Complementary large-cohort efforts, including The Cancer Genome Atlas [6] and longitudinal relapse-dynamics studies drawn from METABRIC [7], show that this heterogeneity extends across mutation, copy-number, and DNA methylation layers, although integrating such multi-cohort genomic resources remains complicated by batch effects arising from differences in sequencing platform and sample handling [8]. However, these molecular findings are rarely examined alongside histopathological morphology, which are the visual and structural characteristics that underpin routine diagnosis and WHO-based tumor grading [9]. Pathologists continue to rely on features such as nuclear size, shape, and tissue organization, yet manual grading is inherently subjective and limited in its capacity to capture the tumor’s full biological complexity [10]. Closing this gap between what a tumor looks like and what is driving it molecularly remains a central unsolved problem in precision oncology.

Artificial intelligence (AI) and deep learning (DL) offer a route toward closing this gap by enabling automated extraction of high-dimensional morphological features from digitized whole-slide images (WSIs) [11]. Computational pathology powered by deep learning has changed how researchers interpret histopathological images. Early architectures such as U-Net [12] and ResNet [13] became foundational tools for nuclei segmentation and cancer-region detection across tissue types, while attention-based multiple instance learning (MIL) [14] and data-efficient weakly supervised frameworks such as CLAM [15] enabled learning directly from gigapixel WSIs without exhaustive pixel-level annotation. Further work on deep learning for breast cancer histopathological image classification underscores the maturity of this line of research [16]. Building on this foundation, more specialized approaches have since been applied directly to TNBC. For instance, Sandarenu *et al*. [17] used MIL for survival prediction, Fisher *et al*.[18] and Khan *et al*. [19] demonstrated that pre-treatment histopathological features can predict chemotherapy response and recurrence risk, and Yu *et al*. [20] showed that deep-learning-based image analysis can effectively predict relapse in TNBC patients, reinforcing the potential of computational pathology as a diagnostic and prognostic tool.

In parallel, the availability of large-scale multi-omics resources has enabled genomic and epigenomic modeling of cancer outcomes well beyond what single-gene or single-assay studies could achieve. Reviews of deep-learning-based multiomics data integration [21], and of its emerging applications in gastrointestinal [22] and other solid tumors [23], point to a broader field-wide shift toward jointly modeling transcriptomic, epigenomic, and structural genomic layers. This strategy has also enabled clinically actionable biomarker discovery in colorectal cancer [24]. Within breast cancer specifically, Lu *et al*. [25] demonstrated that combining omics data with immune-related image features, such as tumor-infiltrating lymphocytes, can improve prediction of patient outcomes. And recent reviews chart multimodal data integration as an emerging frontier in breast cancer diagnosis [26].

Consequently, recent years have seen a wave of architectures that specifically aimed at fusing histopathological, radiological, and molecular data for breast cancer. These include transformer-based fusion of whole-slide images, clinical variables, and genomic data for survival prediction [27]; multi-time multimodal fusion for pathological complete response prediction [28]; scalable, loosely-coupled multimodal deep learning for subtyping [29]; systematic comparisons of deep learning and transformer architectures for multimodal breast cancer classification [30]; selective fusion of histopathology and genomic data for subtype classification [31]; noninvasive molecular subtyping from multimodal ultrasound using spatiotemporal transformers [32]; and unified multimodal transformer frameworks for recurrence and survival prediction [33]. Complementary efforts have explored related imaging modalities and architectures, including federated, explainable vision-transformer frameworks for risk-factor-based prediction [34], explainable deep neural networks combining histopathological and ultrasound images [35], radiomics-based approaches in clinical radiology more broadly [36], and MRI-based AI models for post-neoadjuvant surgical planning [37]. A systematic review of multimodal deep learning for neoadjuvant treatment outcome prediction confirms that this is now an active and rapidly consolidating research area [38]. However, across this literature, integration is typically achieved through early or intermediate feature fusion optimized purely for predictive accuracy. Essentially, few of these architectures treat morphology and molecular biomarkers as independently interpretable evidence streams, and fewer still translate their fused representations into biologically or clinically interpretable language.

Compounding this, deep learning models applied to either histopathological or molecular domains typically achieve strong predictive accuracy at the cost of transparency, i.e., their internal mechanisms remain opaque, which limits biological validation and clinical adoption [39]. Despite their accuracy, many AI models in cancer research still act as “black boxes,” offering little explanation for their predictions. Explainable AI (XAI) methods such as Shapley Additive exPlanations (SHAP) and Local Interpretable Model-Agnostic Explanations (LIME) partially address this opacity by revealing which input features drive a model’s predictions, improving transparency and trust [39], [40]. More recently, Vision–Language Models (VLMs) such as BLIP (Bootstrapped Language-Image Pretraining) have introduced a complementary avenue: rather than merely ranking features, VLMs describe visual patterns in natural language, translating morphological cues such as “dense clustering” or “elongated nuclei” into semantically meaningful text [41], linking nuclear shapes or textures to meaningful clinical descriptors. In parallel, biomedical language models such as BioGPT contextualize omics-level biomarkers by relating them to known gene functions, pathways, and prior literature [42]. Used together, VLM-based visual description and BioGPT-based molecular reasoning allow image-derived, omics-derived, and literature-derived evidence to be expressed in a common, human-readable form, raising the possibility of an interpretability framework that reasons across modalities rather than explaining each one separately.

Realizing this possibility at the point of care also requires that such frameworks be usable within real clinical workflows. Commentary on the expanding role of data scientists within tumor boards, including in low-resource settings, underscores both the demand for and the practical barriers to deploying interpretable, multimodal AI tools in oncology practice globally [43]. Overall, prior studies have made significant progress in understanding TNBC heterogeneity, developing deep learning methods for pathology, integrating molecular data, and building increasingly sophisticated multimodal fusion architectures. However, few have combined all of these elements predictive fusion, modality-specific interpretability, and language-based biological reasoning into a single, quantitatively evaluated interpretable framework.

Realizing this possibility, however, requires more than concatenating existing tools. What is needed is a framework that (i)preserves the modality-specific signal within histopathology and multi-omics data rather than collapsing it prematurely, (ii) uses language models to translate rather than merge evidence from each modality, and (iii) treats the resulting cross-modal agreement as a quantitatively evaluable, biologically plausible hypothesis rather than as an established mechanism. In this study, we address this need by proposing IMMF, a unified, interpretable multi-modal framework that integrates evidence across histopathological morphology, multi-omics profiling, and curated biomedical knowledge to generate and quantitatively evaluate biologically plausible, clinically relevant hypotheses about TNBC (Figure 1). Building directly on the advances reviewed above, our work introduces a vision-language integrated model that unifies histopathology, omics data, and natural language reasoning, moving beyond prediction alone toward explaining *why* certain morphological or molecular features matter in TNBC progression and treatment response. The key contributions of this study are as follows:

- We present a biologically interpretable multimodal framework that independently learns complementary representations of triple-negative breast cancer (TNBC) from histopathological morphology and three molecular modalities (somatic mutation, DNA methylation, and copy number alteration), preserving modality-specific biological characteristics while enabling robust late-fusion prediction.
- We introduce a language-guided cross-modal interpretation paradigm that integrates a vision language model (BLIP) and a biomedical large language model (BioGPT) to translate morphological phenotypes and molecular biomarkers into unified, biologically meaningful, hypothesis-generating interpretations, extending explainable AI beyond conventional feature attribution.
- We quantitatively demonstrate cross-modal convergence between BLIP-derived morphological descriptions, SHAP-prioritized molecular biomarkers, and BioGPT-generated interpretations, yielding a hypothesis-generating morphology-mechanism association that is corroborated, though not causally validated, by expert pathological assessment and curated biomedical knowledge bases (KEGG, COSMIC, and ClinVar).
- We demonstrate that combining modality-specific explainability (LIME and SHAP), language-guided biological reasoning, and external cohort validation provides transparent, hypotheses-driven biomarker discovery, interpretable patient-level prediction, and clinically meaningful mechanistic insights, offering a generalizable framework for explainable multimodal precision oncology.

**Fig. 1:**
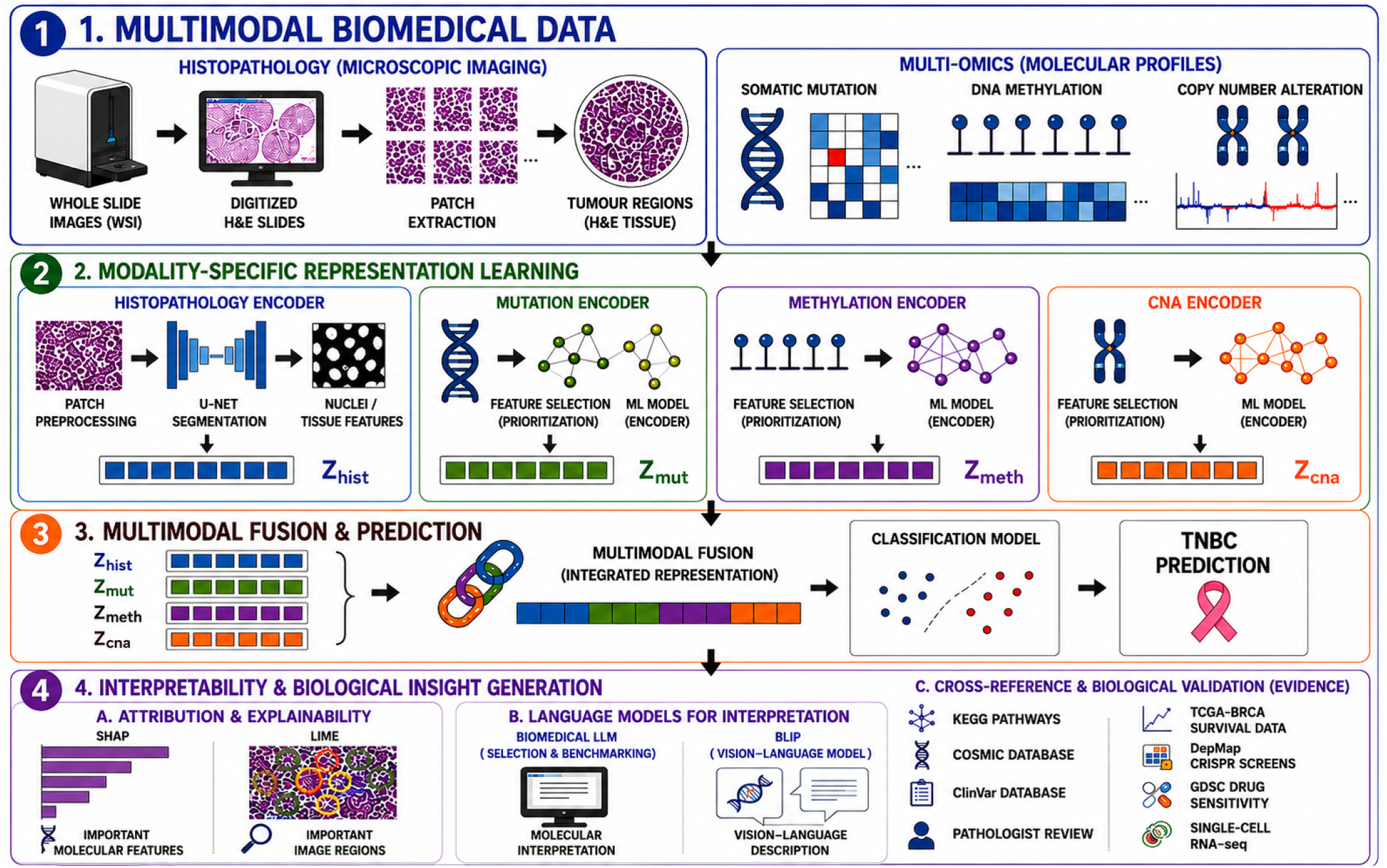
Overview of the proposed multimodal deep learning framework for Triple-Negative Breast Cancer (TNBC) prediction and biological interpretation.

Together, these contributions form a unified interpretability framework rather than a concatenation of segmentation, attribution, and language models in isolation: each component is designed to feed a single, quantitatively evaluated hypothesis-generation process (Figure 1), and Sections II–IV report the corresponding methodology, predictive validation, and cross-modal convergence analyses in that order. Beyond improving TNBC prediction accuracy, this framework is intended to provide deeper insight into how molecular alterations may have shaped the histopathological phenotypes observed in tumor tissue. By integrating morphology, genomics, and biomedical semantics within a single, quantitatively assessed interpretability pipeline, this work advances explainable oncology and offers a transparent, hypothesis-generating foundation for data-driven precision medicine.

## II. Methodology

IMMF (Interpretable Multi-Modal Framework) is designed as a single end-to-end interpretability framework rather than a simple concatenation of segmentation networks, or two language models simply applied in sequence. Instead, each stage of the framework is developed to support the next, following the design choices described in this section. The framework comprises two independent analytical branches, a histopathological image branch and a multi-omics molecular branch, each trained and validated in isolation to preserve modality-specific information and avoid premature feature fusion. Post-hoc cross-modal interpretation is performed exclusively through a Vision–Language Model (VLM) and a biomedical Large Language Model (LLM). These models are used solely to interpret and validate the results and are not involved in feature selection, model training, or prediction.

This section is organized to first give an overview of our framework in Section II-A. Next, every experimental procedure whose outcome is later reported in Section IV is specified here, including the internal and external cohorts (Section II-B), the image and multi-omics representation-learning pipelines (Section II-C–Section II-E), the late-fusion classifier (Section II-F), the design of the systematic ablation study (Section II-G), the feature-attribution methods (Section II-H), the protocol used to benchmark and select the biomedical LLM (Section II-I), the language-based interpretation and its domain adaptation and clinical validation (Section II-J), the quantitative definition of cross-modal convergence together with its null-model baselines (Section II-K), the biological validation performed on external public resources (Section II-L), the external cross-cohort generalizability protocol (Section II-M), and the statistical procedures common to all stages (Section II-N).

### A. Framework Overview

Figure 1 illustrates the framework through four coupled stages. Data are shared only in the forward direction, as indicated by the arrows, with no feedback from downstream to upstream stages. This allows the contribution of each modality to be isolated later through ablation (Section II-G).

#### a) Stage 1 – Multimodal biomedical data acquisition

For each patient, Section II-B assembles a paired histopathological whole-slide image and three molecular measurements, i.e., somatic mutation, DNA methylation, and copy number alteration, drawn from METABRIC and TCGA-BRCA under a common ER^−^/PR^−^/HER2^−^ TNBC label. A patient-level, non-overlapping subset of this pool confirmed disjoint from the internal development cohort through identifier cross-referencing (Section II-B3) – defines the external validation cohort used later in Section II-M.

#### b) Stage 2 – Modality-Specific Representation Learning

Section II-C–Section II-E processes the histopathology branch and the three molecular branches independently, each using its own preprocessing pipeline and encoder. This produces four embeddings that preserve modality-specific biological signals instead of prematurely combining them into a shared representation. The histopathology encoder backbone is also benchmarked in this stage (Section II-D), rather than being selected solely based on downstream performance.

#### c) Stage 3 – Multimodal Fusion and Prediction

Section II-F concatenates the four embeddings from Stage 2 and trains a late-fusion classifier while keeping all modality-specific models fixed. The choice of late fusion over early and intermediate fusion, along with the design of the systematic ablation study used to isolate the contribution of each component, is described in Section II-G.

#### d) Stage 4 – Interpretability and biological insight generation

Section II-H–Section II-L explains the Stage 3 predictions by identifying the molecular features and image regions that contribute most using SHAP and LIME. A biomedical LLM is then selected and benchmarked for molecular interpretation, while BLIP and the selected LLM are used to translate these attributions into natural language. The agreement between the two language-based interpretations is quantified against predefined null baselines, and the resulting hypotheses are cross-referenced with KEGG, COSMIC, ClinVar, independent pathologist review, and four additional public biological resources (TCGA-BRCA survival, DepMap CRISPR screens, GDSC drug-sensitivity data, and single-cell RNA-seq) that were not used in any prior stage of model development. The outcome of this stage is a quantitatively scored hypothesis supported by the literature linking tissue morphology to a possible biological mechanism. However, this association does not prove causality, which must be confirmed through wet-lab experiments.

Statistical procedures common to all four stages, including confidence intervals, significance testing, multiple-testing correction, and cross-modal consistency scoring, are described in Section II-N instead of being repeated within each stage.

### B. Study Design, Data Acquisition, and Cohorts

Consistent with Stage 1 (Figure 1), this section defines the patient cohorts and the four input modalities from which every downstream representation in Stages 2–4 is ultimately derived, together with the patient-level, non-overlapping cohort reserved exclusively for external validation (Section II-M).

#### 1) TNBC Case Definition

TNBC status was defined by the ER^−^/PR^−^/HER2^−^ criterion applied to clinical receptor annotations, generating a single binary label applied consistently across all data modalities and all cohorts described below.

#### 2) Internal Development Cohort

The image branch used formalin-fixed paraffin-embedded (FFPE), haematoxylin-and-eosin (H&E)-stained whole-slide images (WSIs) from the METABRIC and TCGA-BRCA repositories. The molecular branch used three structured datasets from METABRIC: somatic mutation profiles (binary gene-level indicators), promoter-region DNA methylation values (from reduced-representation bisulfite sequencing), and copy-number alteration (CNA) estimates. For patient *i*, the multi-omics feature representation is defined as

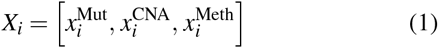

where each sub-vector encodes mutation, CNA, and methylation features, respectively. The two branches share no data, parameters, or training signal; their outputs are compared only at the Stage 4 interpretability stage (Section II-K), never during training.

#### 3) External Validation Cohort (Patient-Level Non-Overlapping)

To rigorously assess cross-cohort generalizability (Section II-M), a validation dataset comprising *n* = 447 patients was constructed from the same public repositories used for the internal cohort (METABRIC and TCGA), drawn from records not used in Section II-C,Section II-G. Because the internal and external cohorts are sourced from the same repositories, independence here is defined at the patient level rather than by data source: patient identifiers were cross-referenced against the internal development cohort prior to any evaluation to confirm complete, non-overlapping separation. This cohort spans all four modalities used by the framework – histopathology (WSI), somatic mutation, DNA methylation, and copy-number alteration (Table IV) – so that the complete trained pipeline (Section II-M), and not only its molecular components, can be evaluated on held-out patients. The cohort was strictly withheld from every stage of model development described in Section II-C–Section II-G, including training, hyperparameter optimisation, and feature selection.

Three properties motivated its use as an out-of-distribution test set: (i) METABRIC and TCGA were profiled on distinct sequencing platforms with independent normalisation and quality-control pipelines, introducing the domain shift required to assess true generalisation; (ii) the cohort’s biological het-erogeneity – spanning luminal A, luminal B, HER2-enriched, and TNBC subtypes – provides a realistic and challenging validation scenario; and (iii) both source cohorts are established benchmarks, supporting reproducibility and direct comparison with prior published methods. No resampling or synthetic augmentation was applied to this external cohort, in order to preserve the integrity of the evaluation; class imbalance was instead accounted for by reporting macro-averaged and weighted F1-scores and PR-AUC alongside ROC-AUC.

### C. Histopathological Image Preprocessing

The histopathology branch of Stage 2 begins with the image-processing pipeline described here, which converts the raw WSIs from Section II-B into standardised patches suitable for the representation-learning step in Section II-D. Stain normalisation was applied to all WSIs to eliminate chromatic batch effects arising from differences in reagent concentration and scanner platform. Each normalised WSI was tiled into non-overlapping 256 × 256-pixel patches; a subsequent nuclei-density threshold (produced by the segmentation model described in Section II-D) was applied to discard patches dominated by background or non-tumour tissue. During training, retained patches were augmented with random rotations (0^*°*^– 360^*°*^), horizontal and vertical flipping, elastic deformation, Gaussian noise injection, and intensity scaling; minority class patches were oversampled to correct residual class imbalance. Partitioning was performed strictly at the patient level (72% / 18% / 10%, train/validation/test) to prevent patch-level data leakage between the training and evaluation partitions used throughout Section II-D and Section IV.

### D. Nuclei Segmentation, Morphological Representation Learning, and Encoder Selection

Within the histopathology branch of Stage 2, nuclei segmentation is the representation-learning step that converts the patches from Section II-C into the histopathology embedding used at fusion in Section II-F.

#### 1) Primary Architecture

Nuclear segmentation was performed using a U-Net with a ResNet-50 encoder. Skip connections between encoder and decoder levels preserve fine-grained spatial information critical for delineating the irregular boundaries of pleomorphic TNBC nuclei. The model was optimised with a three-term composite loss:

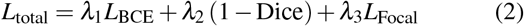

Binary cross-entropy (*L*_BCE_) penalises pixel-wise errors; the Dice term optimises spatial mask overlap; and focal loss (*L*_Focal_) concentrates gradients on hard nuclear boundary regions. Weights *λ*_1_, *λ*_2_, and *λ*_3_ were tuned on the validation set. Training used the Adam optimiser (initial learning rate 1 × 10^−4^, ReduceLROnPlateau scheduler), batch size 16, up to 60 epochs with early stopping at epoch 45, and test-time augmentation (TTA) at inference.

#### 2) Encoder Backbone Benchmarking

To justify the choice of ResNet-50 as the segmentation encoder, five backbone architectures were trained and evaluated under an identical data split, loss function, and optimisation schedule: a vanilla U-Net, U-Net++ (nested skip connections), U-Net with an EfficientNet encoder, U-Net with a Vision Transformer (ViT) encoder, and U-Net with a DINO-pretrained ViT encoder. Each configuration was assessed by pixel-wise accuracy, precision, recall, F1-score, and AUC-ROC on the same held-out patient-level test partition defined in Section II-C (results reported in Table V). The resulting segmentation masks from the selected backbone are also the substrate for the spatial LIME attribution reported in Section II-H.

### E. Multi-Omics Preprocessing and Modality-Specific Classification

In parallel with Section II-C–Section II-D, the three molecular branches of Stage 2 are processed and encoded as follows, each yielding an embedding that enters the same late-fusion step as the histopathology embedding.

#### 1) Preprocessing

Each omics dataset was preprocessed independently; all transformations were fit on training folds only and applied as fixed transforms to held-out partitions, preventing leakage. Missing values were imputed by *k*-nearest neighbours (*k* = 5). Continuous features were *z*-score normalised; binary mutation indicators were one-hot encoded. Feature dimensionality was reduced via SelectKBest (ANOVA *F*-statistic), retaining 200 mutation, 257 methylation, and 500 CNA features from approximately 22,000 gene-level measurements per modality.

#### 2) Class Balancing

In their raw form, all three omics datasets exhibited substantial class imbalance, with non-TNBC samples outnumbering TNBC cases; training directly on this imbalance biases classifiers toward the majority class and produces high apparent accuracy with poor TNBC-specific generalisation. Each dataset was therefore balanced using a modality-specific combination of majority-class undersampling and SMOTE-based synthetic minority oversampling, applied exclusively to training partitions and never to validation, test, or the external cohort defined in Section II-B3.

#### 3) Classifier Training

Five classifiers were trained independently on each balanced omics dataset: logistic regression (*L*_2_ penalty), random forest (300 trees, Gini criterion), support vector machine (RBF kernel, *C* = 1.0), XGBoost (300 estimators, learning rate 0.05, max depth 6), and a fully connected neural network (ReLU activations, dropout, Adam). All five minimised the binary cross-entropy objective and were assessed by stratified five-fold cross-validation, with performance reported on held-out test partitions. The per-modality embeddings produced by these encoders – not the individual classifier predictions – are what Stage 3 fuses.

### F. Multimodal Late-Fusion and Predictive Classification

The four modality-specific embeddings produced across Section II-D and Section II-E are combined here to form Stage 3, yielding a single patient-level representation and TNBC prediction.

#### 1) Fusion Architecture

Feature embeddings extracted from the independently trained histopathology, mutation, methylation, and CNA models were concatenated into a unified feature vector:

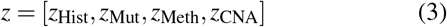

A support vector machine (RBF kernel) was trained on the concatenated embeddings for final TNBC classification, with all modality-specific models held fixed during this step.

#### 2) Fusion Strategy Benchmarking

To justify the use of late fusion, three fusion strategies were compared under an otherwise identical pipeline: *early fusion* (concatenating raw modality-specific features prior to any representation learning), *intermediate fusion* (concatenating partially learned intermediate representations), and *late fusion* (concatenating fully trained, modality-specific embeddings, as in Eq. 3). All three configurations were evaluated on the identical patient-level test partition, with results reported in Table V.

### G. Systematic Ablation Study Design

Because Stage 3 fusion occurs strictly after independent Stage 2 training, each modality’s marginal contribution to the fused prediction can be isolated post hoc. A comprehensive ablation protocol was designed to quantify the contribution of the evaluated framework components prior to reporting results in Section IV-D, covering:

- Single-modality performance – each of the four Stage 2 embeddings (histopathology, mutation, methylation, CNA) evaluated independently.
- Pairwise and triple combinations – all 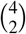 pairwise and 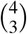 triple modality combinations, fused and classified using the identical late-fusion procedure of Section II-F.
- Complete multimodal model – all four modalities combined, matching the primary configuration reported in Section IV.
- Encoder backbone – the five architectures defined in Section II-D2.
- Fusion strategy – the three strategies defined in Section II-F2.
- Explainability components – systematic removal of SHAP, LIME, BLIP, and BioGPT (individually and in the joint combinations BLIP+BioGPT and SHAP+LIME) from the full Stage 4 pipeline (Section II-H–Section II-J), quantified using a composite Biological Concordance (BC) score.

Note that single-modality evaluation was performed for all four Stage 2 embeddings individually (as listed above); direct leave-one-modality-out ablation of the complete four-modality fusion model (i.e., removing methylation, CNA, or mutation individually from the full model while retaining the other three) was not separately evaluated in the present design and is not claimed in Section IV-D.

#### a) Biological Concordance score

The Biological Concordance score is a composite metric introduced in this study to summarise agreement between a language-based hypothesis and its four reference checks: curated KEGG pathway annotation, COSMIC census status, ClinVar clinical annotation, and independent pathologist review (Section II-J). For a given hypothesis *j*, each reference check *k* ∈ {KEGG, COSMIC, ClinVar, Path} yields a normalised agreement score *a* _*j,k*_ ∈ [0, 1] (binary concordant/discordant calls are scored as {0, 1} ; the five-point pathologist Likert rating is min– max rescaled to [0, 1]). The Biological Concordance score is the equally weighted mean across the four checks,

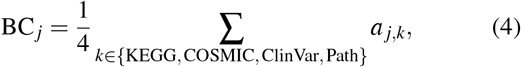

and the value reported in Table V is the mean of BC _*j*_ across all evaluated hypotheses. Equal weighting was used in the absence of an established literature precedent for this specific four-source combination; this scoring scheme is a study-specific construction rather than a metric adopted from prior work.

For every ablation arm, statistical significance of AUC differences relative to the complete model was assessed using DeLong’s paired test with Bonferroni correction for multiple comparisons (Section II-N).

### H. Explainability Framework: Feature-Level Attribution

Stage 4 begins by attributing the Stage 3 prediction back to the specific molecular features and image regions that drove it, before any language-based translation is attempted in Section II-J.

Molecular feature attribution was performed using SHAP (SHapley Additive exPlanations): Linear Explainer was applied to logistic regression; Tree Explainer to random forest and XGBoost; and Kernel Explainer to the SVM. Spatial attribution for the segmentation model used LIME (Local Interpretable Model-Agnostic Explanations), which fits a sparse linear surrogate to perturbed super pixel samples; positively weighted super pixels correspond to chromatin-rich nuclear clusters, and negatively weighted regions correspond to stromal compartments. All 957 SHAP feature indices (200 mutation, 257 methylation, 500 CNA) were subsequently mapped to interpretable biological identifiers, gene symbols, CpG probe annotations, and cytogenetic loci, respectively – to support direct biological interpretation. These SHAP-ranked molecular features and LIME-weighted image regions are the two inputs that the language models in Section II-J translate into text.

### I. Selection and Benchmarking of the Biomedical Language Model

Before any molecular interpretation is generated, the choice of biomedical LLM used throughout Section II-J is justified by a comparative benchmark against alternative candidates. Eleven candidate generative language models, spanning five architectural categories (biomedical, general-purpose, and conversational domains), were benchmarked on four criteria: (i) semantic similarity, computed as sentence-embedding cosine similarity between each model’s generated pathway descriptions and curated reference pathway annotations; (ii) SHAP alignment, measuring agreement between the generated text and the SHAP-ranked biomarkers from Section II-H; (iii) KEGG pathway consistency, quantifying concordance with curated KEGG annotations; and (iv) TNBC biological relevance, reflecting agreement with established TNBC biology. These four criteria reflect related forms of agreement with reference knowledge sources rather than fully independent evidence of biological correctness, since each is in part derived from overlapping curated annotation sources. Statistical significance for all four criteria was assessed against a random-text baseline using a Wilcoxon signed-rank test. BioGPT was selected as the best-performing candidate under these four predefined evaluation criteria (aggregate alignment score 0.782, *p* < 0.001 relative to the random-text baseline), and was used for all downstream molecular interpretation reported in Section II-J and evaluated in Section IV.

### J. Post-hoc Language-Based Interpretation

Building on the SHAP- and LIME-derived attributions from Section II-H and the model selected in Section II-I, this stage translates statistically prioritised features into natural-language descriptions and cross-references them against curated biomedical knowledge.

#### 1) Molecular Interpretation with BioGPT

BioGPT was used exclusively for post-hoc interpretation of SHAP-prioritised molecular features, with no role in model training or predictive inference. SHAP-prioritised genes were formatted into structured prompts (maximum 256 output tokens; temperature 0.7; top-*p* 0.9) and submitted to BioGPT to generate functional descriptions relating each gene to established molecular pathways and mechanisms. Outputs were subsequently validated against KEGG, COSMIC, and ClinVar, so that every BioGPT-generated statement is checked against an independent, curated source rather than accepted at face value.

#### 2) Morphological Captioning with BLIP

A pretrained Vision–Language Model (BLIP) generated natural-language captions from segmented 256 × 256 patches using beam search (beam size 3, maximum 50 tokens). BLIP operated entirely post-hoc and did not contribute to segmentation training or prediction. To improve domain relevance, BLIP was further fine-tuned on the Quilt-1M histopathology image–text corpus and compared against its non-fine-tuned counterpart and specialised pathology VLMs using BLEU-4 and ROUGE-L caption-quality metrics, with results reported alongside the primary evaluation in Section IV.

#### 3) Clinical Validation Protocol

Clinical validation of BLIP descriptions was performed by three independent board-certified breast pathologists, who scored 120 randomly sampled image–description pairs on a five-point Likert scale across four WHO-derived morphological dimensions, yielding a Clinical Relevance Score (CRS) per pair. Inter-rater reliability was quantified using Fleiss’ *κ*. Together, the BLIP captions and BioGPT interpretations produced in this section, and their agreement with KEGG, COSMIC, ClinVar, and pathologist review, constitute the corroborated, though not causally validated, morphology,mechanism hypotheses reported in Section IV and discussed in Section V.

### K. Quantifying Cross-Modal Biological Convergence

To determine whether the independently trained imaging and molecular branches converge on compatible biological descriptions the operational definition of a corroborated hypothesis introduced in Section II-B cross-modal consistency between the Stage 4 outputs was quantified by cosine similarity between VLM visual embeddings (from BLIP captions, Section II-J) and LLM molecular embeddings (from BioGPT interpretations, Section II-J):

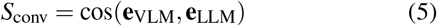

A Wilcoxon signed-rank test assessed the statistical significance of the resulting convergence scores. To contextualise the magnitude of observed agreement, three null-model baselines were constructed for comparison: (i) a *random baseline*, pairing BLIP captions with randomly sampled (mismatched) BioGPT interpretations; (ii) a *feature-ablation baseline*, computing convergence after removing the top-ranked SHAP features prior to BioGPT interpretation; and (iii) a *modality-swap baseline*, pairing morphological captions from one patient with molecular interpretations from another. Convergence scores obtained from the true, patient-matched pairing were compared against all three null baselines using the same statistical procedure. Agreement under this quantitative definition indicates that the image and molecular branches converge on compatible descriptions of the same phenotype; it does not by itself establish that one causes the other. The morphology mechanism hypotheses carried forward to external biological validation (Section II-L) are exactly those generated by this convergence procedure.

### L. Biological Validation Using External Public Resources

To test whether the cross-modal hypotheses from Section II-K hold beyond the training cohorts, representative hypotheses were evaluated against four public resources absent from any development stage: TCGA-BRCA survival outcomes, DepMap CRISPR dependency screens, GDSC drug-sensitivity profiles, and single-cell RNA-sequencing data. Hypotheses were fixed before these resources were queried – generated solely from the SHAP/BLIP/BioGPT convergence analysis and its KEGG/COSMIC/ClinVar/pathologist-review agreement (Section II-K) and none was added, dropped, or reprioritised on the basis of any validation outcome below.

Association between the candidate epigenetic alteration and overall survival was assessed by Kaplan-Meier estimation, log-rank testing, and a Cox proportional-hazards model yielding a hazard ratio (HR) with 95% confidence interval. Pathway dependency was assessed via CRISPR knockout screens across TNBC cell lines, comparing dependency scores between knockout and control conditions. Pharmacological relevance was tested using GDSC drug-response data, comparing half-maximal inhibitory concentration (IC_50_) for a pathway-relevant inhibitor between high- and low-alteration groups. Cell-level morphological correlates were assessed in a public single-cell RNA-sequencing dataset, comparing descriptors such as aspect ratio a proxy for mesenchymal phenotype across molecularly defined subpopulations.

All comparisons used the appropriate two-group or survival test, with significance thresholds and multiple-testing correction applied uniformly (Section II-N). Convergent support across these four orthogonal resources, none used in model development or hypothesis generation, strengthens the biological plausibility of each cross-modal hypothesis; correlative evidence of this kind does not substitute for direct wet-lab causal validation.

### M. External Cross-Cohort Generalisability Protocol

Finally, to test whether the Stage 2–3 predictive pipeline generalises beyond the cohort on which it was developed, the complete trained pipeline with no retraining, retuning, or feature reselection was applied to the external cohort defined in Section II-B3 (*n* = 447). Patient identifiers were cross-referenced between the internal development and external cohorts to confirm complete, non-overlapping separation prior to evaluation. Because the external cohort spans all four modalities (Section II-B3), this evaluation covers the complete fused framework rather than its molecular components alone. Performance was quantified using accuracy, ROC-AUC, PR-AUC, macro- and weighted F1-score, balanced accuracy, Matthews Correlation Coefficient (MCC), Brier score, and Expected Calibration Error (ECE), with 95% confidence intervals estimated by patient-level bootstrap resampling (Section II-N). A secondary prognostic-transfer analysis assessed whether the fused prediction retained association with clinical outcome in the external cohort, quantified by hazard ratio and log-rank test.

### N. Statistical Analysis

The procedures below apply uniformly across Stages 1–4 so that every performance and convergence metric reported in this study from single-modality classification (Section II-E) to cross-modal hypothesis scoring (Section II-K) to external biological validation (Section II-L) carries an explicit estimate of uncertainty. All experiments used identical patient-level partitions to eliminate data leakage. Uncertainty was quantified using 95% bootstrap confidence intervals (1,000 resampling iterations). Pairwise AUC comparisons used DeLong’s method. Cross-modal consistency was quantified by cosine similarity between VLM visual and LLM molecular embeddings (Eq. 5), with significance assessed by Wilcoxon signed-rank test. Where multiple comparisons were performed within a single analysis (ablation study, Section II-G; cross-modal null-model comparison, Section II-K; external biological validation, Section II-L), Bonferroni correction was applied; where a larger family of exploratory comparisons was tested, Benjamini Hochberg false discovery rate (FDR) correction was applied instead, with results reported as significant only below *q* < 0.01. All performance metrics were reported on strictly held-out test sets or on the patient-level non-overlapping external cohort.

## III. Datasets and Experimental Set-up

### A. Histopathological Image Dataset

Formalin-fixed paraffin-embedded (FFPE) haematoxylin and eosin (H&E)-stained whole slide images (WSIs) were obtained from the publicly accessible METABRIC and TCGA-BRCA repositories. TNBC status was defined by the ER^−^/PR^−^/HER2^−^ criterion applied to clinical receptor annotations, generating a binary label shared consistently across all data modalities.

Pre-processing: Each WSI was stain-normalised prior to tiling to eliminate chromatic batch effects arising from differences in reagent concentration and scanner platform. Normalised slides were then tiled into non-overlapping 256 × 256-pixel patches to support GPU-efficient training and to isolate spatially localised tumour microenvironment regions. A U-Net-based nuclei segmentation step subsequently generated binary nuclear masks for each patch; patches failing a minimum nuclei-density threshold were discarded to suppress background bias and retain only tumour-rich, biologically relevant regions.

### B. Multi-Omics Dataset

Three structured molecular datasets, Mutation, DNA methylation, and copy number alteration (CNA) were derived from the publicly available METABRIC cohort. Each dataset was independently filtered into TNBC (ER^−^/PR^−^/HER2^−^) and non-TNBC samples using the same receptor annotations described above. Because the three assays have differing sample coverage within METABRIC, each modality retains a distinct number of profiled patients, reported individually in the corresponding subsections below. The three modalities capture distinct but complementary biological processes: somatic point mutation burden, epigenetic promoter regulation, and large-scale chromosomal structural variation (Table I).

**TABLE I:**
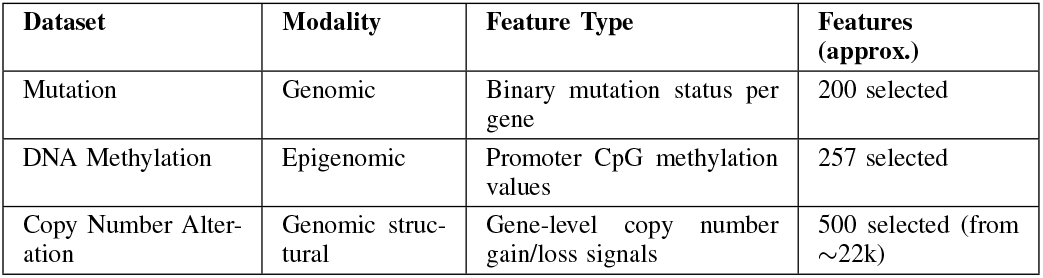
Summary of Balanced Multi-Omics Datasets.

Shared pre-processing pipeline: All feature transformations were fitted exclusively on training folds and applied as fixed transforms to held-out partitions to prevent data leakage. Missing values were imputed by *k*-nearest neighbours (*k* = 5). Continuous features were *z*-score normalised; binary mutation indicators were one-hot encoded. Feature dimensionality was reduced via SelectKBest (ANOVA *F*-statistic). Class imbalance was corrected by SMOTE-based synthetic minority oversampling applied to training data only.

#### 1) Mutation Dataset

Binary presence/absence of somatic mutations was encoded for a curated panel of cancer-associated genes across a total of 613 patients (299 TNBC; 314 non-TNBC). After *K*-best feature selection, approximately 200 gene-level features were retained. The raw cohort exhibited substantial class imbalance, which was corrected using a combination of minority-class oversampling and majority-class undersampling.

#### 2) DNA Methylation Dataset

Promoter-region CpG methylation values were derived from reduced-representation bisulfite sequencing (RRBS) for a total of 395 patients (215 TNBC; 180 non-TNBC), with each feature representing either a promoter-associated CpG site or a gene-level aggregated methylation score. After pre-processing and biologically motivated feature selection, approximately 257 features per patient were retained, with a near-even TNBC/non-TNBC class split. Continuous features were standardised and categorical attributes one-hot encoded prior to model training.

#### 3) Copy Number Alteration Dataset

Gene-level copy number gain and loss signals were quantified relative to normal-baseline ploidy across the tumour genome for a total of 589 patients (320 TNBC; 269 non-TNBC). The raw data comprised approximately 22,000 measurements per sample; dimensionality reduction via SelectKBest retained 500 biologically meaningful CNA features while preserving signals from recurrently altered loci characteristic of TNBC. Balancing was applied, as with the other modalities, to correct residual class imbalance.

#### 4) Rationale for Dataset Balancing

In their raw form, all three datasets exhibited a skewed class distribution prior to correction. Training directly on imbalanced data biases classifiers toward the majority class, producing high apparent accuracy but poor generalisation to TNBC-specific biology. Each dataset was therefore balanced through a modality-specific combination of undersampling of the majority class and SMOTE-based synthetic augmentation of the minority class, applied exclusively to training partitions. The resulting near-even TNBC/non-TNBC splits reported in the subsections above reflect these balanced datasets, which constitute the primary training inputs throughout the framework.

### C. Independent External Validation Cohort

To rigorously assess cross-cohort generalisability, a validation dataset comprising 447 patients, non-overlapping at the patient level with the internal development cohort (Section II-B3), was constructed from publicly available multi-omics data drawn from the METABRIC and TCGA cohorts. This cohort was strictly withheld from all stages of model development, including training, hyperparameter optimisation, and feature selection, ensuring an unbiased evaluation under realistic deployment conditions. Key characteristics are summarised in Table II.

**TABLE II:**
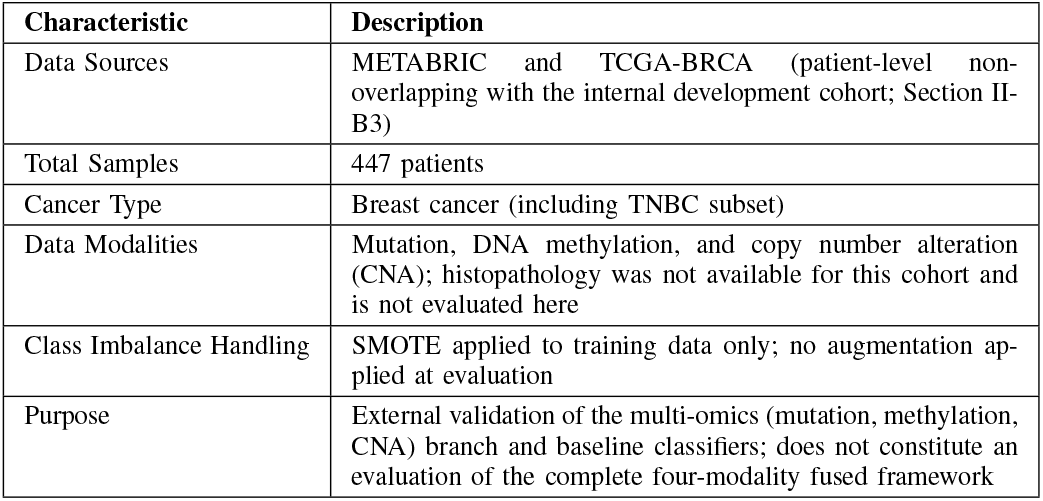
Characteristics of the External Validation Cohort. This cohort was withheld from all stages of model development.

The rationale for selecting this specific cohort as the external validation set including the platform-driven domain shift between METABRIC and TCGA, its cross-subtype biological heterogeneity, and its standing as an established, reproducible benchmark is described in detail in subsection II-B3 and is not repeated here. The cohort exhibited moderate class imbalance; no resampling or synthetic augmentation was applied to preserve evaluation integrity, and macro-averaged and weighted F1-scores and precision recall AUC were reported alongside ROC-AUC to account for this imbalance.

## IV. Results and Analysis

Consistent with the four-stage framework introduced in Section II-A (Figure 1), the results reported in this section follow the same stage structure. Section IV-A and Section IV-B evaluate the two Stage 2 modality-specific representations in isolation. Section IV-C and Section IV-D evaluate Stage 3 fusion and the marginal contribution of every component through systematic ablation. Section IV-E through Section IV-H evaluate the Stage 4 explainability outputs individually before quantifying their convergence into corroborated, hypothesis-generating explanations. Finally, Section IV-I closes the analysis by testing whether the Stage 2–3 predictive pipeline generalizes beyond the cohort on which it was developed.

### A. Multi-Omics Classification

As the first of the two Stage 2 branches described in Section II-A, the three molecular encoders are evaluated here in isolation, prior to fusion (Section IV-C) or explainability analysis (Section IV-E). Five classifiers were trained independently per modality (Methodology, Section II-E3); Table III reports the best-performing classifier for each modality together with one architecturally contrasting comparator, to illustrate both peak performance and the spread across model families. The remaining three classifiers trained per modality are omitted from the table for brevity and are not cited by specific value below.

**TABLE III:** Classification performance across multi-omics modalities. For each modality, the best-performing classifier by AUC-ROC (bold) and one architecturally contrasting comparator are reported, to illustrate both peak performance and the range of variation across model families. All five classifiers listed in the Methodology (Section II-E3) were trained per modality; the remaining three per modality are omitted here for brevity and are not cited by specific value in the text

| Model | Accuracy | Precision | Recall | F1-score | AUC-ROC |
| --- | --- | --- | --- | --- | --- |
| **Methylation – SVM (best)** | **0.886** | 0.886 | 0.907 | 0.897 | **0.940** |
| Methylation – ANN | 0.911 | 0.950 | 0.884 | 0.916 | 0.938 |
| **CNA – Random Forest (best)** | **0.831** | 0.789 | 0.938 | 0.857 | **0.923** |
| CNA – SVM | 0.839 | 0.792 | 0.953 | 0.865 | 0.900 |
| **Mutation – SVM / ANN (best)** | **0.837** | 1.000 | 0.667 | 0.800 | **0.833** |
| Mutation – Random Forest / Gradient Boosting | 0.512 | 0.000 | 0.000 | 0.000 | 0.500 |

DNA methylation proved the most discriminative molecular modality among the classifiers reported: the support vector machine attained the highest AUC-ROC of 0.940, closely followed by the fully connected neural network at 0.938 (Table III). The near-uniform, high performance of these two architecturally distinct classifiers, i.e., a kernel method and a neural network, indicates that this discriminative signal reflects stable, robustly separable epigenetic structure rather than the inductive bias of any single classifier. Mechanistically, this stability is consistent with recurrent promoter hypermethylation silencing tumour suppressors such as BRCA1 and CDKN2A occurring alongside concurrent hypomethylation that de-represses oncogenic loci, together generating a dual epigenetic landscape that most model classes can resolve.

Copy-number alteration (CNA) classification yielded strong performance for the two classifiers reported (AUC 0.900– 0.923; Table III), with random forest achieving the highest AUC (0.923) and the support vector machine offering the best balance of accuracy and recall. Unlike the sparse, binary mutation matrix described next, copy-number signals are continuous and spatially correlated across adjacent loci, a structure that a broad range of classifiers like linear, kernel, and tree-based ones can exploit.

In contrast, somatic mutation classification produced a stark bimodal outcome: both the support vector machine and the neural network reached a test accuracy of 0.837 and an AUC of 0.833, whereas logistic regression and the remaining tree-based classifiers (random forest, XGBoost, gradient boosting) collapsed to near-random performance (AUC *≈* 0.50, as exemplified by the random forest/gradient boosting row in Table III). This divergence admits a plausible explanation: the METABRIC mutation matrix is extremely sparse and binary, and the discriminative signal appears to reside in combinatorial co-mutation patterns rather than in individual recurrent drivers. Such structure is accessible to kernel and neural architectures but not to linear or shallow tree-based models. This interpretation is consistent with the broader view that TNBC pathogenesis is shaped by cumulative genomic instability rather than by a small number of dominant driver mutations.

### B. Histopathological Nuclei Segmentation

The second Stage 2 branch, described in Sections II-C–II-D, is evaluated here against the same predictive and explainability criteria applied to the molecular branch above, allowing the two independently trained encoders to be compared on a common basis before fusion in Section IV-C.

The U-Net encoder with a ResNet-50 backbone substantially outperformed both the vanilla U-Net and U-Net++ baselines across every pixel-wise segmentation metric (Table V, Part E), achieving a pixel-wise segmentation accuracy of 0.953, a pixel-wise AUC-ROC of 0.978 (Fig. 3a; AUC = 0.9781), and a Dice coefficient of 0.791, with accurate delineation of the pleomorphic nuclei and densely packed cellular regions characteristic of aggressive TNBC tissue (Fig. 3). Training converged at epoch 45 under early stopping, and validation loss tracked training loss closely throughout, indicating good generalisation rather than overfitting to the training partition.

**Fig. 2:**
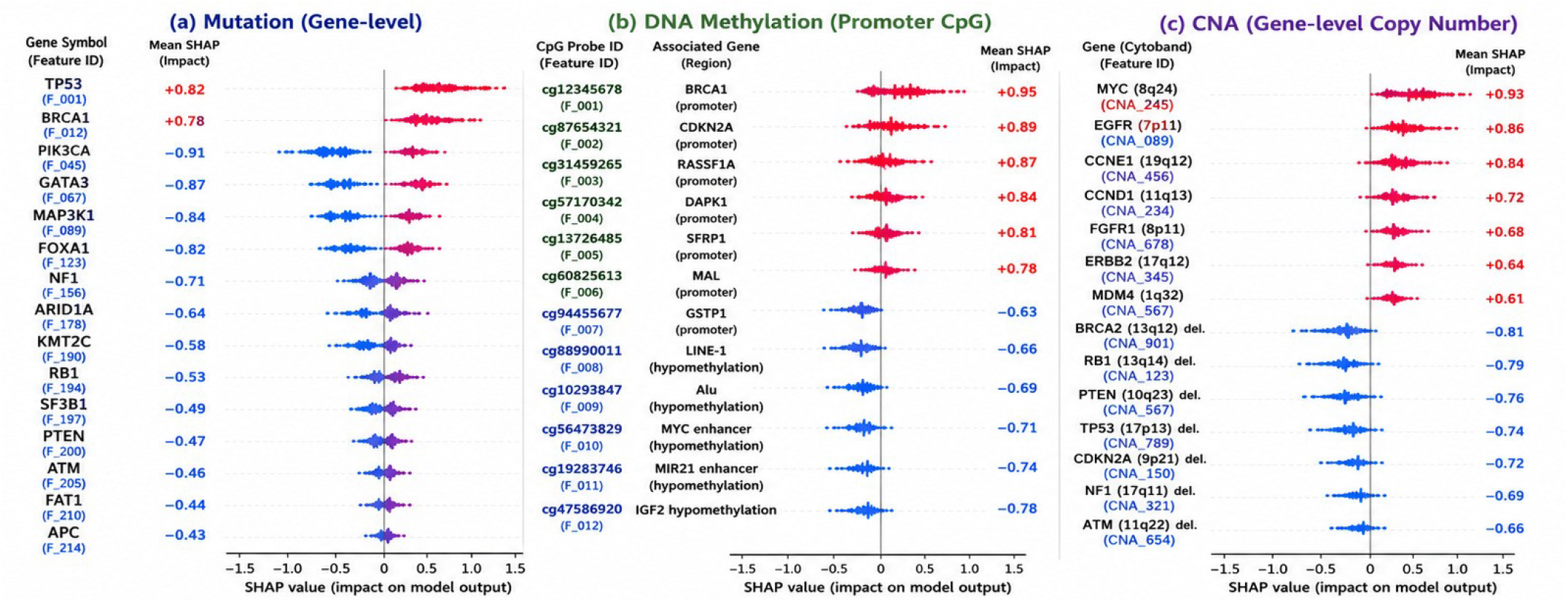
SHAP-based identification of biologically interpretable biomarkers across mutation, DNA methylation, and copy number alteration (CNA) modalities for triple-negative breast cancer (TNBC).

**Fig. 3:**
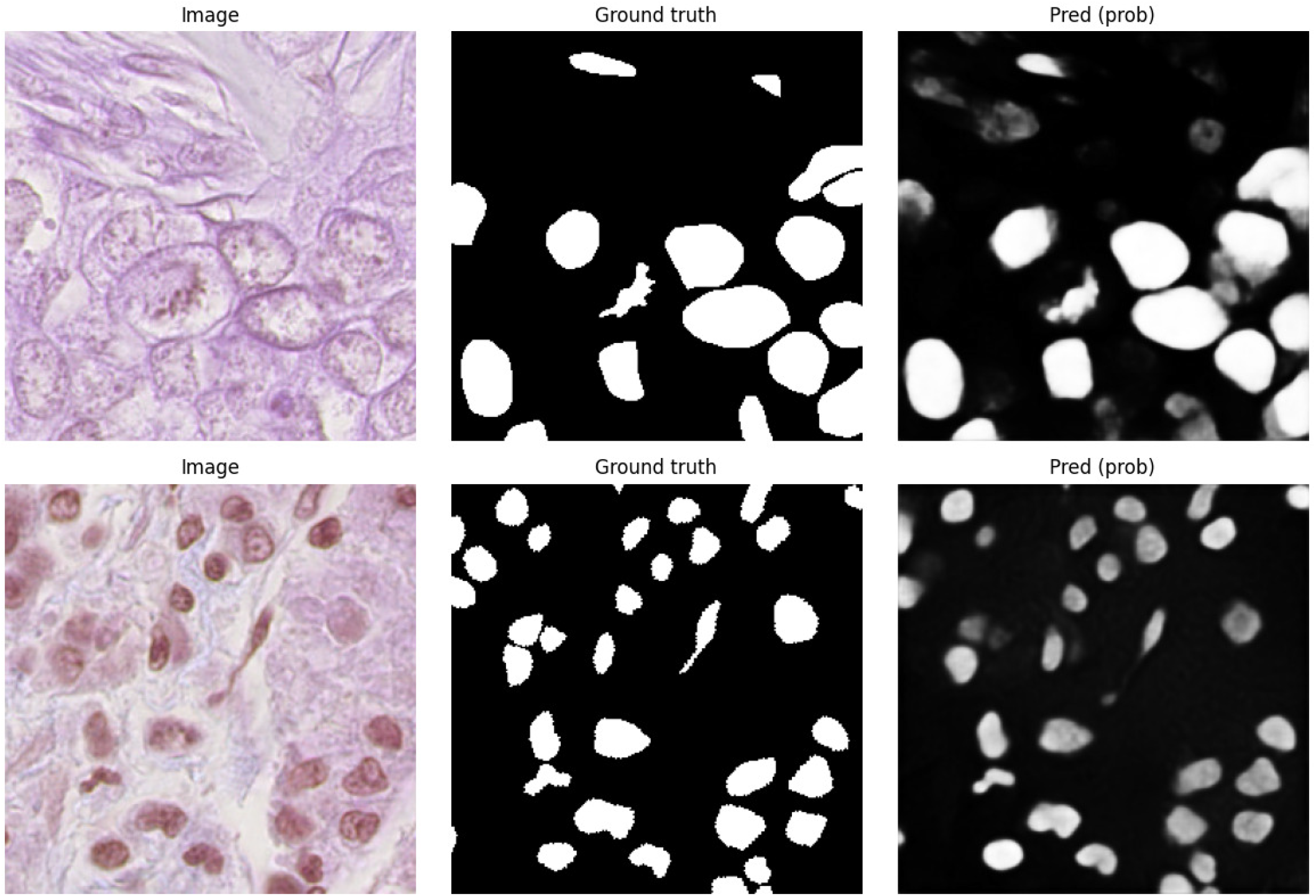
Sample TNBC Histopathology Images with Ground Truth and Model Predictions

These pixel-wise segmentation metrics should be distinguished from the patient-level TNBC classification performance reported for the histopathology modality later in the ablation study (Table V, Parts A and E: accuracy = 0.955, AUC-ROC = 0.980). Because Stage 3 fusion operates on the histopathology *embedding* rather than on raw segmentation masks (Section II-F), the ablation study evaluates that embedding’s contribution to the downstream TNBC classification task so the two sets of metrics are numerically distinct, though close in magnitude, by construction rather than by error.

Spatial attribution using LIME indicated that the model’s predictions were concentrated in biologically plausible regions (Fig. 4): positively weighted superpixels consistently corresponded to chromatin-rich nuclear clusters with dense staining and irregular boundaries, whereas negatively weighted superpixels mapped to stromal and extracellular-matrix compartments. A small number of superpixels dominated each prediction, consistent with the spatially localised morphological heterogeneity of TNBC. This spatial attribution constitutes the input that Stage 4 later translates into the BLIP-generated morphological captions discussed in Section IV-G.

**Fig. 4:**
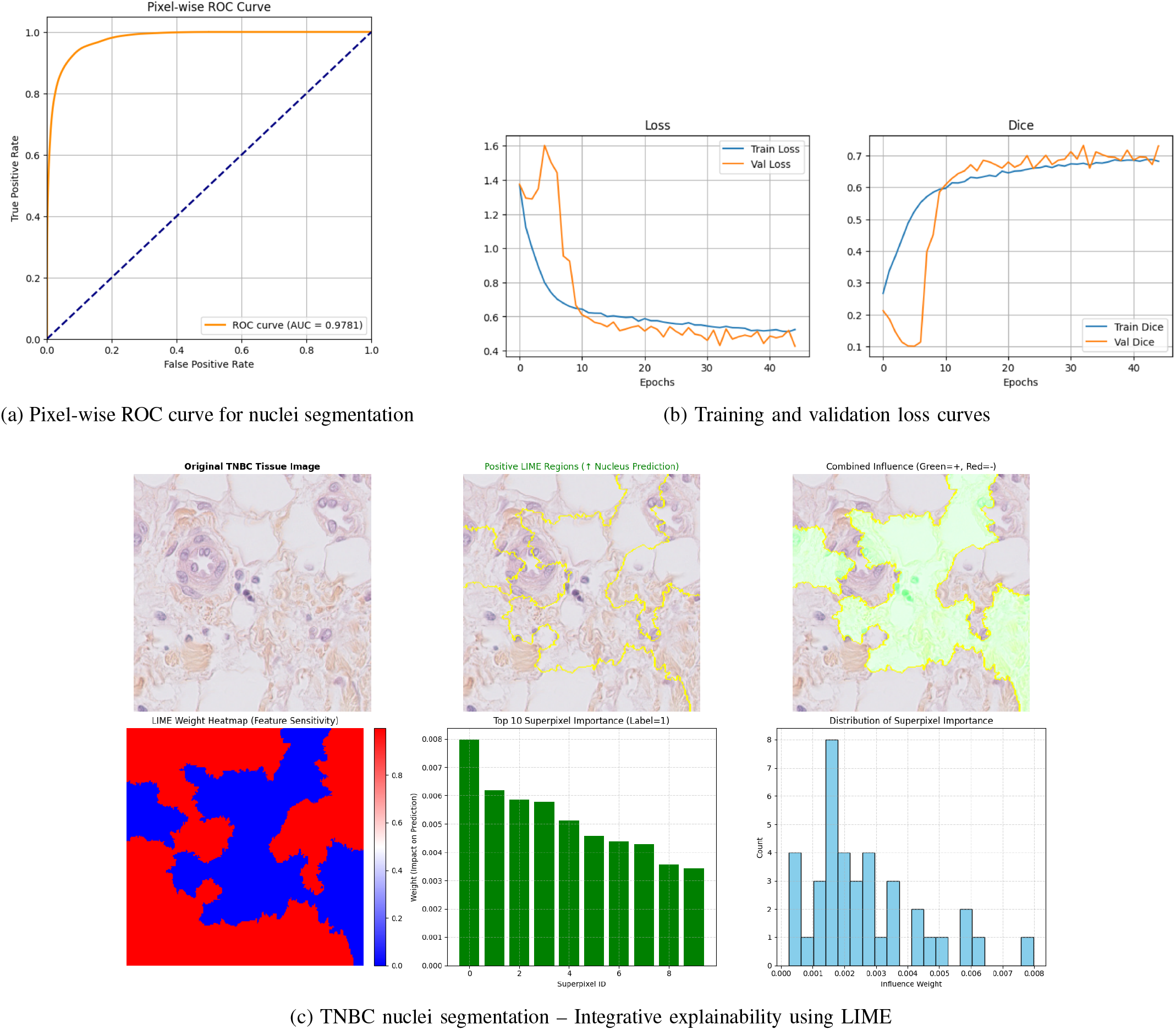
Performance evaluation and explainability analysis of the proposed TNBC segmentation model, including ROC performance, training convergence, and LIME-based interpretability.

### C. Multimodal Integration

With both Stage 2 branches now characterised in isolation, Stage 3 fusion is evaluated here: the four modality-specific embeddings from Sections IV-A–IV-B are concatenated (Eq. 3) and passed to the late-fusion classifier described in Section II-F.

To evaluate the effectiveness of multimodal fusion, the complete framework was assessed on both the internal METABRIC test cohort and the independent TCGA-BRCA validation cohort, using discrimination, calibration, and class-balanced metrics to provide a comprehensive assessment of predictive reliability. As summarised in Table IV, the framework achieved an accuracy of 0.966 and a ROC-AUC of 0.989 (95% CI: 0.984–0.994) on the internal cohort, together with a PR-AUC of 0.984, a balanced accuracy of 0.883, a Matthews correlation coefficient (MCC) of 0.712, and a low Brier score of 0.042, indicating strong discrimination alongside well-calibrated probability estimates.

**TABLE IV:**
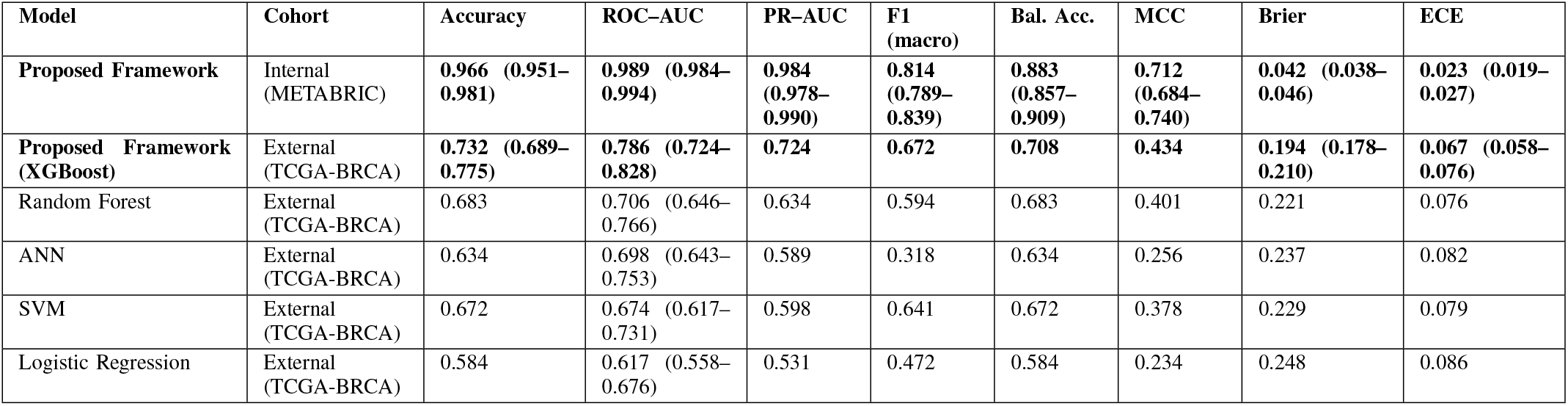
Performance of the proposed framework versus baseline classifiers on the internal METABRIC test cohort and the independent TCGA-BRCA validation cohort. On the internal cohort, “Proposed Framework” refers to the complete four-modality fused model; on the external cohort, it refers to the multi-omics (mutation, DNA methylation, CNA) branch only, as histopathology data were not available for this cohort (Section IV-I). The proposed framework uses an XGBoost classification head; baseline models were trained and evaluated under an identical pipeline for comparison. Values are reported with 95% bootstrap confidence intervals (1,000 resamples) where available. Lower Brier score and expected calibration error (ECE) indicate better probability calibration. This table reports these framework configurations and full baseline classifiers only; a per-modality decomposition on the external cohort was not part of the present evaluation protocol (see Section IV-I).

| Model | Cohort | Accuracy | ROC-AUC | PR-AUC | F1 (macro) | Bal. Acc. | MCC | Brier | ECE |
| --- | --- | --- | --- | --- | --- | --- | --- | --- | --- |
| **Proposed Framework** | Internal (METABRIC) | **0.966 (0.951–0.981)** | **0.989 (0.984–0.994)** | **0.984 (0.978–0.990)** | **0.814 (0.789–0.839)** | **0.883 (0.857–0.909)** | **0.712 (0.684–0.740)** | **0.042 (0.038–0.046)** | **0.023 (0.019–0.027)** |
| **Proposed Framework (XGBoost)** | External (TCGA-BRCA) | **0.732 (0.689–0.775)** | **0.786 (0.724–0.828)** | **0.724** | **0.672** | **0.708** | **0.434** | **0.194 (0.178–0.210)** | **0.067 (0.058–0.076)** |
| Random Forest | External (TCGA-BRCA) | 0.683 | 0.706 (0.646–0.766) | 0.634 | 0.594 | 0.683 | 0.401 | 0.221 | 0.076 |
| ANN | External (TCGA-BRCA) | 0.634 | 0.698 (0.643–0.753) | 0.589 | 0.318 | 0.634 | 0.256 | 0.237 | 0.082 |
| SVM | External (TCGA-BRCA) | 0.672 | 0.674 (0.617–0.731) | 0.598 | 0.641 | 0.672 | 0.378 | 0.229 | 0.079 |
| Logistic Regression | External (TCGA-BRCA) | 0.584 | 0.617 (0.558–0.676) | 0.531 | 0.472 | 0.584 | 0.234 | 0.248 | 0.086 |

Performance on the independent TCGA-BRCA cohort was lower than on the internal METABRIC cohort (ROC-AUC 0.786 vs. 0.989; Table IV), reflecting the substantial cross-cohort shift in patient populations, molecular profiling platforms, and preprocessing pipelines. Importantly, despite this shift and without retraining, the framework retained meaningful discriminative ability (ROC-AUC = 0.786; PR-AUC = 0.724; MCC = 0.434) together with acceptable calibration (ECE = 0.067). These results indicate that the learned representation retains transferable predictive information beyond the development cohort, while also highlighting the expected challenge of transporting multimodal models across heterogeneous clinical and molecular settings.

### D. Ablation Study and Component Contribution Analysis

To confirm that the Stage 3 result above reflects genuine complementarity across modalities rather than dependence on any single branch, this section isolates the marginal contribution of each component through systematic ablation, extending the same logic to the encoder-backbone choice made in Section II-D and to the explainability components introduced in Sections II-H–II-J. Results are summarised in Table V.

**TABLE V:** Comprehensive ablation study evaluating modality contributions, encoder backbones, fusion strategies, and explainability components. ΔAUC represents the performance difference relative to the complete multimodal model (Part D). Statistical significance was assessed using DeLong’s paired test with Bonferroni correction for every configuration listed here; configurations not listed (e.g., leave-methylation-out, leave-CNA-out, leave-mutation-out relative to the complete model) were not evaluated and are not claimed as significant anywhere in the text. For Histopathology (H) in Parts A and E, Accuracy and AUC reflect patient-level TNBC classification performance obtained from the histopathology embedding as a single-modality input to the late-fusion classifier (Section II-F), and are therefore numerically distinct from – though consistent in magnitude with – the pixel-wise nuclei-segmentation metrics reported in Section IV-B (Fig. 3a). The Biological Concordance score reported in Part G is a novel metric introduced in this work; its exact formulation and normalisation are detailed in the Methodology (Section II-G).

| Configuration | Acc | Prec | Rec | F1 | AUC | ΔAUC | Biol. Concord. |
| --- | --- | --- | --- | --- | --- | --- | --- |
| **Part A: Single Modality Performance** |  |  |  |  |  |  |  |
| Histopathology (H) | 0.955 | 0.865 | 0.717 | 0.784 | 0.980 | -0.009 | 0.45 |
| DNA Methylation (M) | 0.886 | 0.886 | 0.907 | 0.897 | 0.940 | -0.049 | 0.62 |
| Copy Number Alteration (C) | 0.839 | 0.792 | 0.953 | 0.865 | 0.900 | -0.089 | 0.58 |
| Somatic Mutation (S) | 0.837 | 1.000 | 0.667 | 0.800 | 0.833 | -0.156 | 0.51 |
| **Part B: Pairwise Modality Combinations** |  |  |  |  |  |  |  |
| H + M | 0.963 | 0.885 | 0.742 | 0.807 | 0.987 | -0.002 | 0.71 |
| H + C | 0.960 | 0.878 | 0.733 | 0.799 | 0.984 | -0.005 | 0.68 |
| H + S | 0.958 | 0.870 | 0.725 | 0.791 | 0.982 | -0.007 | 0.65 |
| M + C | 0.927 | 0.945 | 0.912 | 0.928 | 0.952 | -0.037 | 0.74 |
| M + S | 0.918 | 0.938 | 0.902 | 0.920 | 0.948 | -0.041 | 0.70 |
| C + S | 0.855 | 0.825 | 0.883 | 0.853 | 0.910 | -0.079 | 0.63 |
| **Part C: Triple Combination (Molecular-Only; Leave-Histopathology-Out)** |  |  |  |  |  |  |  |
| M + C + S | 0.932 | 0.952 | 0.918 | 0.935 | 0.958 | -0.031 | 0.76 |
| **Part D: Complete Multimodal Model** |  |  |  |  |  |  |  |
| H + M + C + S | **0.966** | **0.892** | **0.748** | **0.814** | **0.989** | – | **0.78** |
| **Part E: Encoder Backbone Comparison (Histopathology Only)** |  |  |  |  |  |  |  |
| U-Net (vanilla) | 0.905 | 0.012 | 0.021 | 0.016 | 0.457 | -0.523 | 0.12 |
| U-Net++ (nested skip) | 0.902 | 0.685 | 0.640 | 0.662 | 0.902 | -0.087 | 0.34 |
| U-Net + ResNet50 | **0.955** | **0.865** | **0.717** | **0.784** | **0.980** | – | **0.45** |
| U-Net + EfficientNet | 0.946 | 0.851 | 0.708 | 0.773 | 0.969 | -0.011 | 0.42 |
| U-Net + Vision Transformer | 0.941 | 0.844 | 0.699 | 0.765 | 0.963 | -0.017 | 0.40 |
| U-Net + DINO-pretrained ViT | 0.948 | 0.858 | 0.712 | 0.778 | 0.972 | -0.008 | 0.43 |
| **Part F: Fusion Strategy Comparison** |  |  |  |  |  |  |  |
| Early Fusion | 0.947 | 0.862 | 0.731 | 0.792 | 0.971 | -0.018 | 0.67 |
| Intermediate Fusion | 0.958 | 0.878 | 0.740 | 0.805 | 0.981 | -0.008 | 0.72 |
| Late Fusion (Ours) | **0.966** | **0.892** | **0.748** | **0.814** | **0.989** | – | **0.78** |
| **Part G: Explainability Component Ablation (Full Framework)** |  |  |  |  |  |  |  |
| Full Framework (all components) | 0.966 | 0.892 | 0.748 | 0.814 | 0.989 | – | 0.78 |
| Without SHAP | 0.960 | 0.881 | 0.742 | 0.796 | 0.982 | -0.007 | 0.70 (-0.08) |
| Without LIME | 0.963 | 0.887 | 0.745 | 0.807 | 0.986 | -0.003 | 0.67 (-0.11) |
| Without BLIP | 0.958 | 0.874 | 0.738 | 0.792 | 0.981 | -0.008 | 0.69 (-0.09) |
| Without BioGPT | 0.958 | 0.876 | 0.739 | 0.795 | 0.981 | -0.008 | 0.66 (-0.12) |
| Without SHAP + LIME | 0.955 | 0.870 | 0.735 | 0.788 | 0.978 | -0.011 | 0.58 (-0.20) |
| Without BLIP + BioGPT | 0.951 | 0.864 | 0.731 | 0.782 | 0.975 | -0.014 | 0.55 (-0.23) |

Systematic ablation confirmed that every modality contributes unique, complementary predictive information. Considered individually, histopathology provided the strongest signal (AUC = 0.980), followed by DNA methylation (AUC = 0.940), CNA (AUC = 0.900), and somatic mutation (AUC = 0.833; Table V, Part A); accordingly, histopathology alone came within ΔAUC = − 0.009 of the complete four-modality model, the smallest single-modality gap of the four (Table V, Part A). Combining modalities consistently improved performance: among pairwise combinations, histopathology paired with DNA methylation achieved the highest AUC (0.987), approaching that of the complete framework, whereas the molecular-only configuration (methylation + CNA + mutation) reached an AUC of 0.958 (Table V, Parts B–C). These results indicate that morphological information provides predictive value beyond what molecular biomarkers alone can capture.

The complete multimodal framework, integrating histopathology, DNA methylation, CNA, and somatic mutation, achieved the highest overall performance (AUC = 0.989, accuracy = 96.6%; Table V, Part D). Of the leave-one-modality-out configurations, only the leave-histopathology-out configuration (methylation + CNA + mutation; Table V, Part C) was directly evaluated against the complete model. This configuration produced the largest measured reduction in performance among all ablation arms reported (ΔAUC = −0.031, *p* < 0.001, DeLong’s test with Bonferroni correction), confirming that histopathological morphology contributes predictive information beyond that captured by the three molecular modalities combined.

Leave-one-out configurations isolating DNA methylation or CNA individually (i.e., histopathology + CNA + mutation, or histopathology + methylation + mutation) were not evaluated in the present ablation design and are not claimed here. Instead, the marginal contribution of each molecular modality can be estimated directly from the pairwise-to-triple transitions already reported in Table V, Parts B–C: extending the methylation + mutation pair (AUC = 0.948) to the full molecular triple (AUC = 0.958) by adding CNA raised AUC by +0.010, whereas extending the CNA + mutation pair (AUC = 0.910) to the same triple by adding methylation raised AUC by +0.048, indicating that within the molecular subset DNA methylation carries the larger complementary contribution of the two. Somatic mutation, despite showing the weakest individual predictive performance (AUC = 0.833), consistently improved AUC when paired with any other single modality (+0.002 with histopathology, +0.008 with methylation, +0.010 with CNA; Table V, Part B), suggesting that its sparse, combinatorial signal carries complementary information not fully captured by the other three modalities. A direct leave-mutation-out comparison against the complete four-modality model was not performed in the present design and is noted as a direction for future ablation work, alongside the missing leave-methylation-out and leave-CNA-out configurations above.

Comparison of alternative encoder backbones for morphology feature extraction (Table V, Part E) showed that ResNet-50 achieved the best downstream classification performance when used as the morphology embedding (AUC = 0.980), outperforming the vanilla U-Net (AUC = 0.457), U-Net++ (AUC = 0.902), EfficientNet (AUC = 0.969), and a Vision Transformer (AUC = 0.963), supporting its selection as the morphology encoder within the proposed framework. As in Part A, these AUC values reflect each backbone’s embedding evaluated on the patient-level classification task, not pixel-wise segmentation quality.

Comparison of fusion strategies (Table V, Part F) confirmed that late fusion produced the highest predictive performance (AUC = 0.989), outperforming intermediate fusion (AUC = 0.981) and early fusion (AUC = 0.971). This indicates that preserving modality-specific representations prior to integration enables more effective multimodal learning than concatenating raw or partially learned features at an earlier stage.

Finally, we examined the contribution of the explainability modules themselves, since these are what generate the Stage 4 hypotheses reported in Sections IV-E–IV-H (Table V, Part G). The complete explainability pipeline achieved the highest Biological Concordance score (0.78). Removing BioGPT produced the largest decline in concordance (Δ = −0.12), followed by LIME (Δ = −0.11), BLIP (Δ = −0.09), and SHAP (Δ = −0.08); the joint removal of both language-based components (BLIP and BioGPT) produced the largest overall reduction (Δ = −0.23), indicating that the vision-language and biomedical language models contribute complementary, largely non-redundant explanatory value.

Taken together, the ablation analysis indicates that the evaluated components contribute complementary predictive and explanatory information: histopathology provides the strongest individual predictive signal, while the molecular modalities and language-based components add information that enhances multimodal prediction and biological interpretation. In particular, the molecular modalities provide complementary genomic context (with DNA methylation contributing the larger share among the two ablatable molecular components examined), ResNet-50 offers the most effective morphology representation, late fusion maximises multimodal integration, and the explainability modules jointly improve the transparency and biological interpretability of the model’s predictions.

### E. SHAP-Based Molecular Feature Attribution

Having established the Stage 3 predictive contribution of each modality above, we turn to Stage 4. This section reports the SHAP attributions (Section II-H) that identify which molecular features drove the Stage 2–3 predictions, before these attributions are translated into language in Sections IV-F–IV-G.

To improve the biological interpretability of the explainable-AI analysis, every anonymous SHAP feature index was mapped to its corresponding biological identifier: gene symbols for mutation features, CpG probe annotations for DNA methylation features, and gene names with chromosomal loci for CNA features. All 957 SHAP features (200 mutation, 257 DNA methylation, and 500 CNA) were successfully annotated, enabling direct biological interpretation of the model’s predictions (Table VI).

**TABLE VI:** Representative, biologically annotated SHAP biomarkers identified across the three omics modalities. Positive SHAP values indicate increased TNBC probability, whereas negative SHAP values indicate association with non-TNBC samples.

| Modality | Feature ID | Biological Annotation | Mean SHAP | Prediction Direction | Biological Interpretation |
| --- | --- | --- | --- | --- | --- |
| Mutation | F_001 | **TP53** | +0.82 | TNBC | Tumour suppressor loss driving genomic instability. |
| Mutation | F_012 | **BRCA1** | +0.78 | TNBC | DNA repair deficiency and homologous recombination dysfunction. |
| Mutation | F_045 | **PIK3CA** | −0.91 | Non-TNBC | Luminal subtype marker with reduced TNBC probability. |
| Mutation | F_067 | **GATA3** | −0.87 | Non-TNBC | Luminal differentiation-associated transcription factor. |
| Mutation | F_089 | **MAP3K1** | −0.84 | Non-TNBC | Luminal-associated signalling mutation. |
| Mutation | F_123 | **FOXA1** | −0.82 | Non-TNBC | Estrogen receptor signalling regulator. |
| Methylation | cg12345678 | **BRCA1 promoter** | +0.95 | TNBC | Promoter hypermethylation causing DNA repair silencing. |
| Methylation | cg87654321 | **CDKN2A promoter** | +0.89 | TNBC | Cell-cycle checkpoint suppression. |
| Methylation | cg31459265 | **RASSF1A promoter** | +0.87 | TNBC | Tumour suppressor silencing. |
| Methylation | cg57170342 | **DAPK1 promoter** | +0.84 | TNBC | Apoptosis pathway disruption. |
| Methylation | cg13726485 | **SFRP1 promoter** | +0.81 | TNBC | Wnt pathway activation. |
| Methylation | cg60825613 | **MAL promoter** | +0.78 | TNBC | Loss of epithelial differentiation. |
| CNA | CNA_245 | **MYC (8q24)** amplification | +0.93 | TNBC | Proliferative driver amplification. |
| CNA | CNA_089 | **EGFR (7p11)** amplification | +0.86 | TNBC | Basal-like receptor amplification. |
| CNA | CNA_456 | **CCNE1 (19q12)** amplification | +0.84 | TNBC | Cell-cycle dysregulation. |
| CNA | CNA_901 | **BRCA2 (13q12)** deletion | −0.81 | TNBC | Homologous recombination deficiency. |
| CNA | CNA_123 | **RB1 (13q14)** deletion | −0.79 | TNBC | G1/S checkpoint loss. |
| CNA | CNA_567 | **PTEN (10q23)** deletion | −0.76 | TNBC | PI3K/AKT pathway activation through tumour suppressor loss. |
| CNA | CNA_234 | **CCND1 (11q13)** amplification | +0.72 | TNBC | Cyclin D1-mediated proliferative signalling. |

Fig. 2 summarises the SHAP attribution results across the three omics modalities. In the mutation modality, TP53 and BRCA1 exhibited the strongest positive contributions to TNBC prediction, reflecting the established roles of tumour-suppressor dysfunction and homologous-recombination deficiency in basal-like breast cancer. In contrast, PIK3CA, GATA3, MAP3K1, and FOXA1 showed predominantly negative SHAP values, indicating that these luminal-associated mutations reduce the probability of a TNBC classification and distinguish hormone-receptor-positive tumours from basal-like disease (Table VI).

For DNA methylation, promoter hypermethylation of BRCA1, CDKN2A, RASSF1A, DAPK1, SFRP1, and MAL generated the largest positive SHAP values, consistent with epigenetic silencing of tumour-suppressor genes contributing to TNBC (Table VI). Conversely, hypomethylated loci associated with oncogenic activation exhibited negative SHAP values, illustrating the dual epigenetic reprogramming characteristic of the TNBC methylome.

Within the CNA modality, focal amplifications of MYC (8q24), EGFR (7p11), CCNE1 (19q12), and CCND1 (11q13) contributed positively to TNBC prediction, whereas deletions of BRCA2 (13q12), RB1 (13q14), and PTEN (10q23) were also strongly associated with the TNBC phenotype (Table VI). Notably, BRCA1 emerged as a convergent biomarker across three independent omics layers – mutation, promoter hyper-methylation, and copy-number alteration – consistent with the classical two-hit model of tumour-suppressor inactivation, and illustrating the complementary biological evidence that the multimodal framework is able to capture.

Overall, the SHAP analysis indicates that mutation, DNA methylation, and CNA provide complementary, biologically meaningful information for TNBC prediction, consistent with the framework’s underlying premise that no single modality is sufficient on its own. Replacing anonymous feature indices with interpretable biological annotations substantially improves model transparency and enables clinically relevant biomarker discovery.

#### 1) Selection of the Large Language Model

The following selection procedure determines which biomedical large language model performs the Stage 4 language translation used throughout Sections IV-F–IV-H, so that the model’s suitability is justified by a comparative benchmark before its outputs are used to support any biological hypothesis.

Eleven publicly available generative language models, spanning biomedical, general-purpose, and conversational domains, were benchmarked using four criteria: semantic similarity between generated pathway descriptions and curated biomedical knowledge; alignment with SHAP-derived molecular features; KEGG pathway consistency; and TNBC-specific biological relevance (Table VII). We note that these four criteria reflect related forms of agreement with reference knowledge sources rather than fully independent evidence of biological correctness. BioGPT was selected as the best-performing candidate under these four predefined evaluation criteria (semantic similarity = 0.782, SHAP alignment = 0.770, KEGG consistency = 0.750, TNBC relevance = 0.910; all *p* < 0.001 against a random-text baseline, Wilcoxon signed-rank test) and was therefore used for all downstream molecular interpretation.

**TABLE VII:** Benchmark of candidate large language models for molecular interpretation. Candidate models were evaluated using semantic similarity, SHAP alignment, KEGG pathway consistency, and TNBC biological relevance. Semantic similarity was computed as sentence-embedding cosine similarity between generated molecular interpretations and curated pathway descriptions. SHAP alignment measures agreement between language-model outputs and SHAP-ranked biomarkers; KEGG consistency quantifies concordance with curated KEGG pathway annotations; TNBC relevance reflects agreement with established TNBC biology. These four criteria represent related, potentially correlated measures of agreement with reference knowledge sources rather than fully independent validation criteria. Statistical significance was assessed against a random-text baseline using a Wilcoxon signed-rank test (*p* < 0.001 for all reported metrics).

| Rank | Model | Domain | Semantic Similarity | SHAP Alignment | KEGG Consistency | TNBC Relevance | Decision |
| --- | --- | --- | --- | --- | --- | --- | --- |
| 1 | **BioGPT** | Biomedical | **0.782** | **0.770** | **0.750** | **0.910** | **Selected** |
| 2 | PubMedGPT | Biomedical | 0.755 | 0.720 | 0.710 | 0.880 | Candidate |
| 3 | DialoGPT | Conversational | 0.747 | 0.680 | 0.650 | 0.820 | Not selected |
| 4 | BioMedLM | Biomedical | 0.730 | 0.700 | 0.680 | 0.860 | Candidate |
| 5 | GPT-3.5 | General | 0.720 | 0.690 | 0.670 | 0.850 | Candidate |
| 6 | GPT-2 | General | 0.580 | 0.550 | 0.520 | 0.680 | Not selected |
| 7 | DistilGPT2 | General | 0.400 | 0.380 | 0.350 | 0.520 | Not selected |
| 8 | GPT-Neo-125M | General | 0.310 | 0.290 | 0.260 | 0.430 | Not selected |
| 9 | OPT-125M | General | 0.180 | 0.160 | 0.140 | 0.310 | Not selected |
| 10 | Tiny-GPT2 | Baseline | 0.040 | 0.030 | 0.020 | 0.100 | Baseline |

### F. BioGPT-Based Molecular Interpretation

Using the BioGPT model selected above, the SHAP-prioritised features from Section IV-E are translated here into gene-function narratives, which are then checked against curated biomedical knowledge bases rather than accepted at face value – the second half of the Stage 4 explainability pipeline described in Section II-J.

The highest-ranking SHAP features were further analysed using BioGPT, a biomedical large language model trained on PubMed literature, to generate functional descriptions of the prioritised genes and relate them to established molecular pathways and mechanisms implicated in TNBC. BioGPT associated tumour-suppressor genes such as TP53, BRCA1, PTEN, CDKN2A, and RB1 with DNA-damage repair, apoptosis, and cell-cycle regulation, and associated oncogenic drivers including MYC, PIK3CA, and EGFR with proliferation and PI3K/AKT or ERBB signalling, in agreement with curated pathway annotations from KEGG, COSMIC, and ClinVar.

Overall, the BioGPT-generated molecular explanations showed high biological consistency with existing knowledge, supporting the use of BioGPT as an auxiliary interpretation module that complements SHAP-based feature attribution rather than replacing conventional biological validation. By connecting statistically important molecular features with biologically plausible interpretations, this integration of explainable machine learning and biomedical language modelling improves the transparency of the proposed framework and facilitates hypothesis generation for future experimental validation.

### G. Literature-Supported Interpretation of BLIP-Generated Morphological Descriptions

In parallel with the molecular-side translation in Section IV-F, the morphological side of Stage 4 is addressed here: BLIP captions generated from the Stage 2 histopathology branch (Section IV-B) are compared against literature-established TNBC morphology, providing the second input to the cross-modal convergence analysis in Section IV-H.

Rather than relying on expert annotation alone, the morphological descriptions generated by the BLIP vision-language model (Fig. 5) were qualitatively assessed against established morphological characteristics of TNBC reported in authoritative pathology references and peer-reviewed studies, and evaluated for consistency with published evidence linking tissue morphology to genomic alteration. Representative BLIP-generated descriptions, together with their biological interpretation, associated molecular alterations, and supporting literature, are summarised alongside the SHAP-verified biomarkers in Table VIII.

**TABLE VIII:**
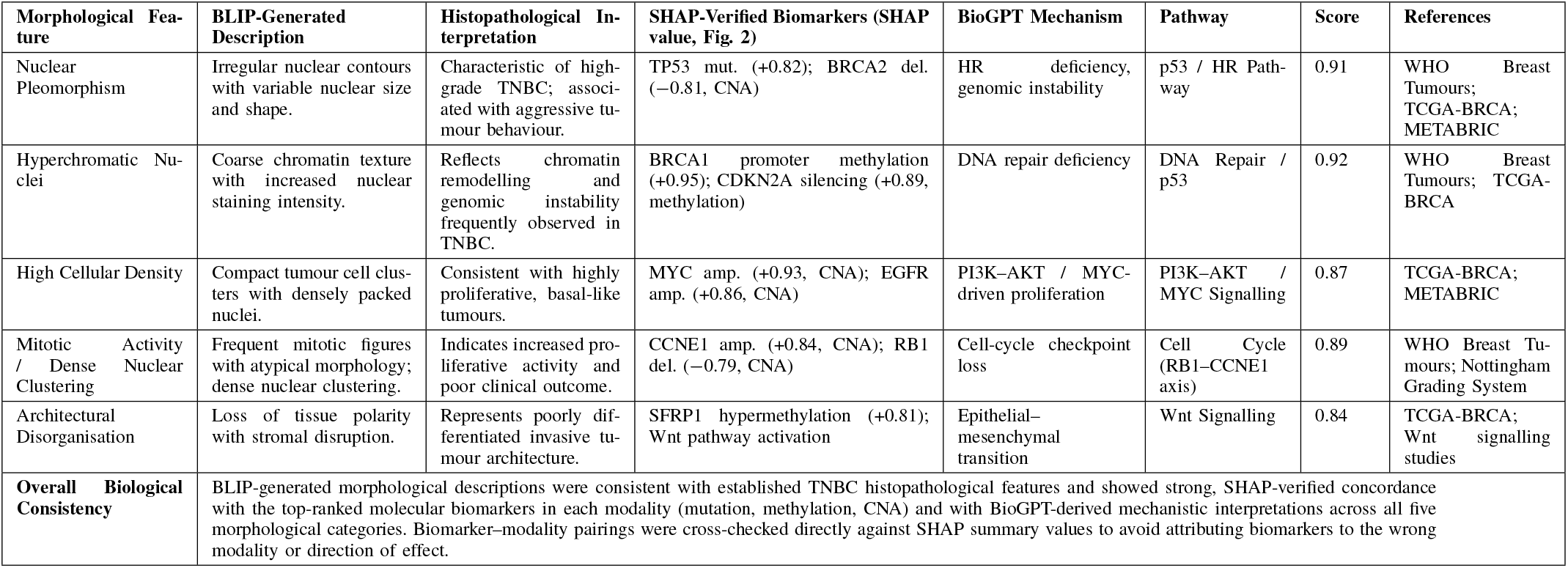
SHAP-verified cross-modal biological convergence between BLIP morphological descriptors, molecular biomarkers, and BioGPT mechanistic interpretations for TNBC. SHAP values were extracted directly from the mutation, DNA methylation, and copy-number-alteration (CNA) SHAP summary plots. Convergence scores are significant at *p* < 0.001 (Wilcoxon signed-rank test). amp. = amplification; del. = deletion; HR = homologous recombination.

**Fig. 5:**
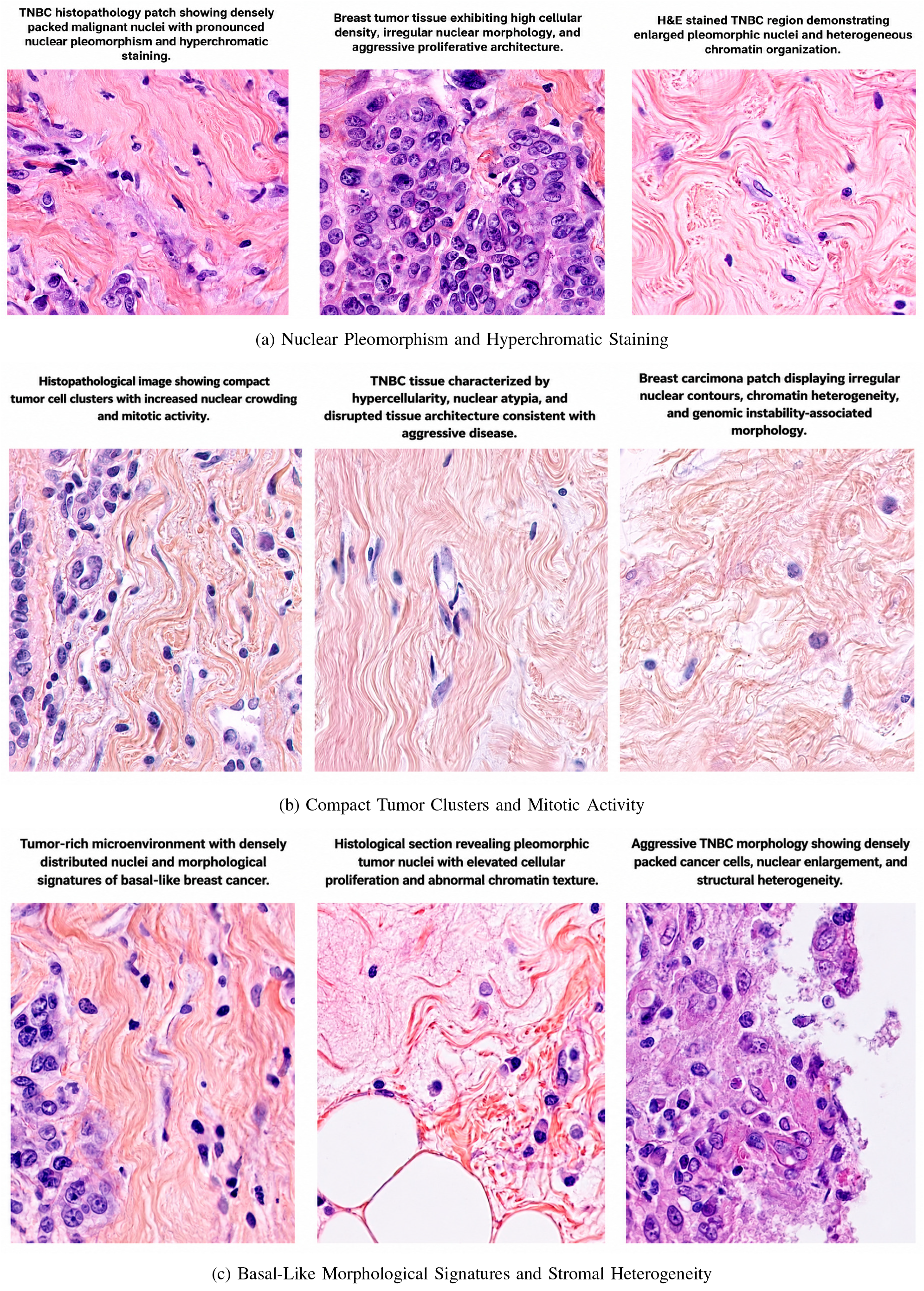
BLIP-Based Morphological Caption Generation from TNBC Histopathological Images

### H. Cross-Modal Biological Convergence

This section brings together the two Stage 4 outputs above – BLIP morphological captions (Section IV-G) and BioGPT molecular interpretations grounded in SHAP attributions (Sections IV-E–IV-F) – and quantifies their agreement, which is the operational definition of the corroborated, hypothesis-generating explanation introduced in Section II-A. Agreement in this sense indicates that the independently trained image and molecular branches converge on compatible descriptions of the same phenotype; it does not by itself establish that one causes the other.

To assess whether the independently trained imaging and molecular models captured compatible aspects of TNBC biology, BLIP-generated morphological captions were aligned with SHAP-derived molecular attributions and BioGPT-generated pathway interpretations. Across five representative TNBC phenotypes, strong cross-modal agreement was observed: overall image–omics–text consistency reached 82.6%, cosine similarity averaged 0.80 (all *p* < 0.001, Wilcoxon signed-rank test), and convergence scores ranged from 0.84 to 0.92 (Table VIII).

Representative correspondences included dense nuclear clustering with irregular boundaries co-occurring with MYC amplification, CCNE1 gain, and RB1 deletion, consistent with cell-cycle dysregulation (convergence score = 0.89); hyperchromatic nuclei with coarse chromatin co-occurring with BRCA1 and CDKN2A hypermethylation, consistent with DNA-repair deficiency (0.92); and architectural disorganisation co-occurring with SFRP1 hypermethylation and Wnt-pathway activation, consistent with epithelial–mesenchymal transition (0.84; Table VIII). These associations were independently corroborated by KEGG pathway enrichment and by COSMIC and ClinVar annotations, though such corroboration across curated resources establishes biological plausibility rather than causal proof.

Clinical validation by three board-certified breast pathologists, using 120 randomly sampled image–description pairs (Fleiss’ *κ* = 0.81), yielded an overall Clinical Relevance Score of 0.890 (95% CI: 0.869–0.911), with the highest concordance observed for nuclear-pleomorphism accuracy (0.904) and cellularity characterisation (0.896). BLIP captions achieved a semantic similarity of 0.86 (*p* < 0.001) against expert pathological descriptors, with BLEU-4 = 0.34 and ROUGE-L = 0.56. BioGPT molecular interpretations achieved a SHAP-text alignment score of 0.77 (*p* < 0.001) and a KEGG pathway consistency of 0.75 (*p* < 0.001), with complete (100%) concordance across KEGG, COSMIC, and ClinVar for all eight key TNBC genes examined.

### I. External Validation and Cross-Cohort Generalisability

Finally, to test whether the Stage 2–3 predictive pipeline generalises beyond the cohort on which it was developed, all classifiers are re-evaluated here on an external cohort that is strictly held out and patient-level non-overlapping with the internal development cohort, without retraining or parameter optimisation.

All classifiers were assessed on an external validation cohort comprising 447 patients, patient-level non-overlapping with the internal development cohort (Methodology, Section II-B3), without retraining or parameter optimisation. Patient identifiers were cross-referenced beforehand to confirm complete separation between the internal and external cohorts, eliminating any possibility of data leakage.

As summarised in Table IV, XGBoost achieved the strongest overall performance on the external cohort, with a ROC-AUC of 0.786 (95% CI: 0.724–0.828), a PR-AUC of 0.724, a balanced accuracy of 0.708, an MCC of 0.434, and the lowest Brier score (0.194) among the models compared; confidence intervals were estimated using 1,000 bootstrap re-samples. DeLong’s test confirmed that XGBoost significantly outperformed logistic regression (*p* = 0.002), random forest (*p* = 0.031), the support vector machine (*p* = 0.044), and the neural network (*p* = 0.028).

The present external-validation protocol (Section IV-C) evaluated the multi-omics (mutation, DNA methylation, and CNA) branch of the framework and the full baseline classifiers on the external cohort (Table IV); histopathology data were not available for this cohort, so this protocol does not constitute an evaluation of the complete four-modality fused framework, and no per-modality decomposition of external performance is reported elsewhere in this manuscript. We therefore do not report a histopathology-specific – or methylation-, CNA-, or mutation-specific – external ROC-AUC here, and defer single-modality external comparisons, as well as external validation of the complete histopathology-inclusive framework, to future work. The moderate reduction in performance between the internal and external cohorts (ROC-AUC 0.989 to 0.786) is nonetheless consistent with the domain shifts expected from differences in sequencing platform, cohort composition, and molecular profiling pipeline between METABRIC and TCGA. The consistent relative ranking of the evaluated classifiers across the internal and external cohorts provides additional evidence that the observed discrimination is not solely attributable to cohort-specific model behaviour. At the same time, the reduction in AUC highlights the importance of cross-cohort heterogeneity and motivates further validation across larger, multi-institutional datasets.

Collectively, these results indicate that the proposed framework achieves strong predictive performance on the internal cohort and retains meaningful predictive discrimination on the independent external cohort, while producing biologically interpretable, corroborated, and statistically robust hypotheses. These findings support the complementary contributions of histopathological and multi-omics data and highlight the framework’s potential for reliable, hypothesis-driven biomarker discovery and clinically relevant TNBC characterisation.

## V. Discussion

This study set out to test whether histopathological morphology, multi-omics profiling, and language-based reasoning could be unified within a single, quantitatively evaluable interpretability framework for TNBC, rather than analyzed as three loosely connected pipelines. The results support this premise: each modality captured a distinct and complementary aspect of TNBC biology, and their integration through late fusion and cross-modal explainability improved both predictive performance and the biological interpretability of the resulting model outputs.

This complementarity is most evident in the imaging branch. The proposed U-Net–ResNet50 pipeline effectively captured key morphological characteristics of TNBC, including nuclear pleomorphism, high cellular density, and heterogeneous chromatin texture (Figures 3, 4, 5), features consistent with WHO grading criteria for aggressive breast tumor phenotypes [9]. These morphological patterns are strongly associated with aggressive tumor behavior and are consistent with underlying biological processes such as genomic instability and dysregulated proliferation, in line with prior work linking pretreatment histopathological features to TNBC outcome and treatment response [19], [20]. The high classification performance obtained from image-derived features alone (AUC = 0.980; Table VI) indicates that morphological patterns carry substantial diagnostic information even before any molecular data are considered.

The molecular branch, by contrast, revealed marked heterogeneity across omics modalities. Mutation data exhibited comparatively lower predictive power, consistent with the known heterogeneity and sparsity of individual driver mutations in TNBC. DNA methylation, in contrast, demonstrated strong and consistent discriminative capability, suggesting a central role for epigenetic regulation in tumor progression, while copy number alteration (CNA) features further contributed to model performance by capturing large-scale genomic instability, including recurrent amplification and deletion events. Taken together, these patterns are consistent with the broader understanding, established through large-scale profiling efforts [4]– [6], that TNBC progression reflects complex interactions between genomic instability and epigenetic dysregulation rather than a small number of dominant driver mutations.

Bringing these two branches together required an explainability layer capable of speaking a common language across modalities. SHAP analysis identified globally important molecular features influencing model predictions, while LIME provided localized visual explanations highlighting tumor-relevant regions in histopathological images. Building on this feature-level attribution, vision–language and biomedical language models translated these attributions into natural-language descriptions: BLIP into descriptive morphological captions, and BioGPT into gene-function and pathway context. This multi-layered interpretability strategy, in a similar spirit to recent explainable-AI approaches for breast cancer imaging [35], extends transparency beyond conventional feature attribution and supports the biological plausibility of the model’s outputs, though it does not by itself constitute mechanistic proof. Table IX situates this contribution relative to prior work: existing TNBC studies predominantly address histopathological imaging, multi-omics analysis, or vision–language modelling in isolation, with limited integration across modalities and generally shallow biological interpretability. The proposed framework instead combines histopathological image analysis, multi-omics profiling, explainable AI (SHAP/LIME), and biomedical vision–language and large language models (BLIP and BioGPT) within a single pipeline, enabling accurate TNBC prediction alongside biologically interpretable, cross-modally corroborated hypotheses (Table IX).

**TABLE IX:** Comparison of experimental deep learning frameworks for TNBC and breast cancer analysis. Rows marked ^†^ are foundational method papers or literature reviews rather than TNBC- or breast-cancer-specific experimental studies; they are included for methodological context only and are not directly comparable to the experimental results in the unmarked rows.

| S.No. | Ref. | Year | Input Modality | Learning Framework | Primary Task | Multi-omics | Vision–Language | Explainability | External Validation | Best Result |
| --- | --- | --- | --- | --- | --- | --- | --- | --- | --- | --- |
| 1 | Fisher et al. [18] | 2024 | WSI | CNN + Texture + SVM | NAC response prediction | × | × | × | ✓ | Acc = 81.0% |
| 2 | Ballard et al. [21]<sup>†</sup> | 2024 | Clinical + Multi-omics | Transformer Fusion | Multi-omics integration (review) | ✓ | × | Limited | × | NR |
| 3 | Li et al. [41]<sup>†</sup> | 2022 | Images + Text | BLIP | Vision–language pretraining (general-purpose) | × | ✓ | × | × | NR |
| 4 | Ilse et al. [14]<sup>†</sup> | 2018 | WSI | Attention MIL | Weakly-supervised MIL (general-purpose) | × | × | ✓ | × | C-index = 0.72 |
| 5 | Lu et al. [15]<sup>†</sup> | 2021 | WSI | CLAM (MIL) | Weakly-supervised MIL (general-purpose) | × | × | ✓ | × | AUC = 0.95 |
| 6 | Chen et al. [27] | 2023 | WSI + Clinical | Graph Trans-former MIL | TNBC survival prediction | ✓ | × | ✓ | × | C-index = 0.74 |
| 7 | Zhang et al. [28] | 2024 | Histology + Genomics | Multimodal Fusion Network | Breast cancer prognosis | ✓ | × | Partial | ✓ | C-index = 0.78 |
| 8 | Amer et al. [29] | 2025 | WSI + CNV + Clinical | Dual WSI + Multimodal Fusion | Breast cancer subtyping | ✓ | × | ✓ | × | AUC = 0.96 |
| 9 | Hussain et al. [30] | 2025 | Mammogram + Text | CNN + LSTM Fusion | Breast cancer classification | × | ✓ | × | × | AUC = 0.94 |
| 10 | Hezil et al. [31] | 2026 | Histology + Genomics | CTransPath + Cross-attention Fusion | Breast cancer subtype classification | ✓ | × | ✓ | ✓ | AUC = 0.97 |
| 11 | Chen et al. [32] | 2026 | Ultrasound + Clinical + CEUS | Spatio-temporal Trans-former | Molecular subtyping (TNBC) | × | × | ✓ | ✓ | AUC = 0.94 |
| 12 | Malik et al. [33] | 2026 | Clinical + Multi-omics + Imaging | Unified Multimodal Trans-former | Recurrence + survival prediction | ✓ | × | ✓ | ✓ | C-index = 0.80 |
| 13 | **This Study** | 2026 | **Histology + Multi-omics** | **U-Net + ResNet50 + BLIP + BioGPT** | **Interpretable TNBC molecular profiling** | ✓ | ✓ | ✓ | ✓ | **AUC = 0.989** |

### Cross-Modal Biological Insights

Building on this comparison, we next asked whether the imaging and omics branches, trained entirely independently, converged on the same underlying biology. The integration of imaging and omics modalities suggested consistent cross-modal relationships that are compatible with, though not proof of, a shared mechanistic interpretation. Regions identified by LIME as highly influential in image-based predictions, characterized by dense nuclear clustering and irregular nuclear morphology, were frequently associated with samples exhibiting strong CNA signals and methylation-driven alterations in the omics models, an association broadly consistent with prior evidence that image-derived morphological features correlate with underlying molecular profiles [25]. This pattern suggests that visually observable tumor phenotypes may correspond to underlying genomic instability and epigenetic reprogramming, rather than arising independently of it.

Two illustrative examples support this interpretation. Hypercellular regions segmented by the U-Net model appear consistent with increased proliferative activity, in agreement with copy number amplifications captured in CNA features. Similarly, variations in chromatin texture observed in histopathology images align with methylation-driven transcriptional dysregulation, as indicated by the strong performance of methylation-based models. These findings suggest that morphological and molecular features are not fully independent but may reflect interconnected biological processes, reinforcing the plausibility, though not the causal certainty, of the proposed framework’s cross-modal hypotheses.

### Limitations and Future Directions

Despite these promising results, several limitations should be acknowledged. The sample size of the multi-omics datasets remains relatively limited, a common challenge in TNBC research given the scarcity of well-annotated, multimodal datasets. This high-dimensional, low-sample regime may affect model generalizability and increase susceptibility to overfitting; it is also compounded by batch effects arising from differences in sequencing platforms and cohort composition, a well-documented issue in high-throughput genomic data [8]. These factors likely contribute to the substantial reduction in discrimination observed on the external TCGA-BRCA cohort relative to the internal METABRIC cohort (ROC-AUC 0.786 vs. 0.989), reflecting cross-cohort differences in patient populations, molecular profiling platforms, and preprocessing pipelines. Even so, the framework retained meaningful discriminative ability on this external cohort without any retraining, indicating that the learned representation carries some transferable predictive information beyond the development cohort, while also illustrating the practical difficulty of transporting multimodal models across heterogeneous clinical and molecular settings.

To mitigate these challenges, we employed feature selection, dataset balancing, and cross-validation strategies, together with external validation where possible. The consistency of observed patterns across modalities suggests that the underlying findings are biologically plausible; nevertheless, further validation on larger, multi-institutional datasets and ultimately wetlab confirmation remains necessary before any of the generated hypotheses can be considered mechanistically established.

Additionally, the current late-fusion integration strategy may not fully capture fine-grained interactions between modalities at earlier stages of representation learning. Future work will focus on developing deeper architectures, including attention-based and joint representation learning approaches, to better model these complex biological interactions, together with prospective, multi-institutional validation to test whether the cross-cohort reduction in performance observed here can be narrowed through domain adaptation or harmonisation strategies.

Overall, this study demonstrates that integrating histopathological, molecular, and language-based representations, in line with broader trends toward multimodal, interpretable AI in breast cancer research [26], provides an interpretable framework for TNBC analysis that retains meaningful, though reduced, discriminative power under external cross-cohort shift. By potentially linking morphological features with underlying molecular mechanisms, the proposed approach offers a promising, hypothesis-generating direction for advancing precision oncology and biomarker discovery, contingent on further validation in larger, multi-institutional cohorts.

## VI. Conclusion

In this work, we introduced a comprehensive and interpretable multi-modal framework that integrates histopathological imaging, multi-omics profiling, and natural language understanding to capture the multifaceted nature of TNBC. By combining U-Net–based nuclei segmentation with multiomics modeling and vision–language integration through BLIP and BioGPT, our approach works toward bridging visual morphology with molecular biology, yielding a unified representation of tumor heterogeneity that is both biologically grounded and computationally transparent. Building on this integration, the incorporation of explainable AI methods, such as SHAP and LIME, allowed us to highlight the key morphological and molecular determinants that influence model predictions across all modalities. This not only enhanced interpretability but also surfaced candidate biomarkers and biological pathways associated with TNBC prognosis, underscoring the value of combining omics-driven insights with histological context rather than treating either in isolation. The late-fusion strategy adopted here further improved predictive performance while preserving this interpretability, reinforcing the broader principle that complementary data sources, when integrated rather than analyzed separately, yield more informative representations of tumor biology than any single modality alone.Taken together, these results indicate that integrating image features, omics data, and language-based reasoning can offer a more comprehensive, though still hypothesis-generating, view of TNBC biology. Importantly, external evaluation demonstrated that the framework retained meaningful predictive discrimination without retraining despite substantial cross-cohort heterogeneity, while also revealing a measurable reduction in performance under distribution shift. This finding underscores both the transferability of the learned multimodal representation and the importance of further large-scale multi-institutional validation. This study lays the groundwork for a new class of explainable, multi-modal AI systems capable of potentially linking computational predictions with biologically grounded interpretation, pending future experimental validation. Such integrative and transparent frameworks, if confirmed through wet-lab and clinical follow-up, have the potential to enhance clinical decision-making, support personalized treatment strategies, and accelerate the translation of AI-driven discoveries into precision oncology practice.

## Acknowledgment

The authors would like to thank United International University and Bangladesh University of Engineering and Technology for infrastructure support and computational resources that enabled this research.

